# Learning precise spatiotemporal sequences via biophysically realistic circuits with modular structure

**DOI:** 10.1101/2020.04.17.046862

**Authors:** I. Cone, H. Z. Shouval

## Abstract

The ability to express and learn temporal sequences is an essential part of learning and memory. Learned temporal sequences are expressed in multiple brain regions and as such there may be common design in the circuits that mediate it. This work proposes a substrate for such representations, via a biophysically realistic network model that can robustly learn and recall discrete sequences of variable order and duration. The model consists of a network of spiking leaky-integrate-and-fire model neurons placed in a modular architecture designed to resemble cortical microcolumns. Learning is performed via a learning rule with “eligibility traces”, which hold a history of synaptic activity before being converted into changes in synaptic strength upon neuromodulator activation. Before training, the network responds to incoming stimuli, and contains no memory of any particular sequence. After training, presentation of only the first element in that sequence is sufficient for the network to recall an entire learned representation of the sequence. An extended version of the model also demonstrates the ability to successfully learn and recall non-Markovian sequences. This model provides a possible framework for biologically realistic sequence learning and memory, and is in agreement with recent experimental results, which have shown sequence dependent plasticity in sensory cortex.

## Introduction

So long as time flows in one direction, nature itself is fundamentally sequential. To operate in this reality, the brain needs to think, plan, and take action in a temporally ordered fashion. When you sing a song, hit a baseball, or even utter a word, you are engaging in sequential activity. More accurately, you are engaging in sequential recall of a learned activity – your actions not only have *a* temporal order and duration, but *the* temporal order and duration which you observed. Hence, the question of how sequence representations are learned, stored, and recalled is of fundamental importance to neuroscience. Recent evidence has shown that such learned representations can exist in cortical circuits^1–5^, begging the question: how are these circuits constructed, and how can such representations be learned?

To address these questions, we introduce a biophysically realistic modular network with an eligibility trace-based learning rule that can robustly learn and recall both the order and duration of elements in a sequence. Although the model’s construction is based upon recent experimental observations in visual cortex^1–3^, utilizing observed cell types^6–8^ and laminar structure^9,10^, many of its key aspects (modularity, heterogenous representations) are illustrative of general principles of sequence learning. The ability of the network to learn both duration and order, along with its use of a learning rule that bypasses the need for constant and explicit targets, differs from most historical and contemporary models of sequence learning^11–20^. We also present an extended formulation of the model, which is capable of learning and recalling sequences with non-Markovian (i.e. history dependent) transitions.

A variety of different models have been proposed to account for the representation and learning of sequences. Most of these models fall into one of two classes: chain structures or recurrent neural networks (RNNs). Chain structure models of neural sequence learning operate in a method akin to synfire chains – that is, representations of different individual stimuli are linked together, in order, via feed forward synaptic connections^11–13^. However, this method alone is insufficient for capturing the full functionality of sequence representation and learning. In such a chain-like structure, there is nothing encoding start times, stop times, or durations of the individual elements of the sequence. In general, such chain structures can learn and encode the order of sequence elements, but not other temporal properties of the sequence. Some models have attempted to address these issues via ad-hoc solutions such as variable adaptation time constants^15^, but these typically require the network to have a priori information about the sequence it will be representing.

Recurrent neural nets (RNNs) are another common class of sequence learning models^17–20^. Unlike explicit feed forward chain models, RNNs learn sequences by leveraging rich, dynamical representations to approximate target outputs. However, common learning rules for these models, such as backpropagation through time (BPTT) or FORCE are biologically unrealistic, as they require some combination of non-local information, precise and symmetrical feedback structures, and/or explicit feedback about the targets at every time point^21^. Interestingly, recent work has shown that networks can under some conditions learn inputs via random feedback connections rather than backpropagation^22^, but this random feedback is less effective than BPTT for learning sequences with long timescales^23^.

Our model takes elements from both chain and recurrent models in order to establish a new, hybrid framework for sequence learning which avoids the pitfalls of either method when used in isolation. The model’s modular structure enables it to learn both the order and duration of sequence elements, and to do this with biophysically plausible learning rules.

## Results

### A Model for Sequence Learning Based on Modular Architecture and Eligibility Trace Learning

In order to learn a sequence, a network needs to learn both the order of elements as well as the duration of each element. In a traditional “chain-like” network with Hebbian learning, order of presented stimuli can be readily encoded by learning directional feed forward connections between populations. However, this simple architecture proves insufficient for representing the duration of presented stimuli and their specific start and end times, as intrinsic time constants determine the speed at which the signal travels through the network^14,15^.

This issue is illustrated in Figure 1. In this schematic model, each module responds to a specific external stimulus. An external sequence activates these different modules in a given order and with externally determined durations for each element (Fig. 1a). Learning can change the feedforward synaptic efficacies between modules to reflect the order of the presented stimuli. Upon recall, triggered by activating the first stimulus, each of the of the encoded stimuli are activated at the correct order. However, the duration of activation within the module, as well as the timing of the activation of the subsequent module, are determined by the intrinsic time constants of the network (Fig. 1b), causing a temporal compression of the recalled sequence. While certain time constants, asymmetrical inhibition, or adaptation with slow time constants could be manually placed in the network to facilitate recall of the particular sequence in Figure 1a, there does not yet exist a biophysically realistic, robust, and general solution to this problem.

**Figure 1.**
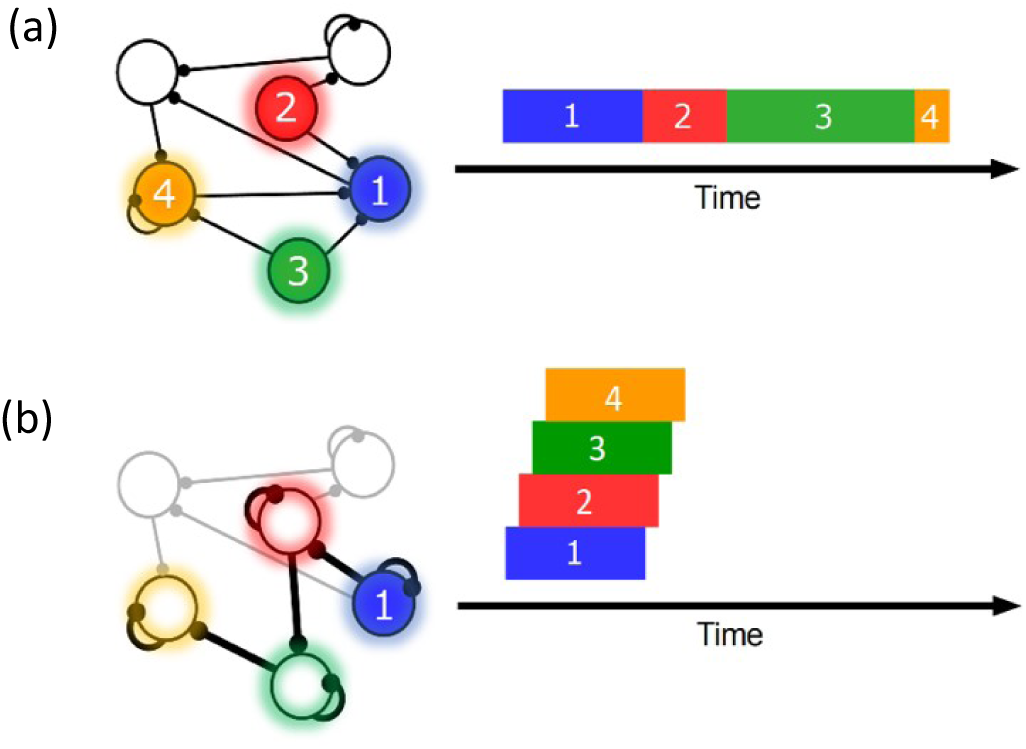
Sequence representation in networks. **a)** A network composed of different populations of cells, each population is activated by a specific stimulus, and there are plastic connections between and within these populations. Initially these connections are random and weak. Upon presentation of a sequence of stimuli (filled circles, left), the populations will become activated for the duration and in the order in which they are stimulated (right). **b)** After many presentations of a particular sequence, successful asymmetric Hebbian learning encodes the order of the stimuli into the synaptic weights of the network. After training, upon presentation of the first element of the sequence (filled circle, left), the network can recall (right) the order of presentation, but the timing and duration of each element is lost. In a generic network such as this, the timing of recall is determined by intrinsic time constants of the system and not the duration in the sequence that was presented.

In this work we present a network of spiking neurons that can learn both the order and duration of sequence elements. This model uses biophysically realistic learning rules, in contrast to most RNN models^17–20^. As in the “chain-like” models, the order is learned by modifying the feed-forward synaptic efficacies between modules. Unlike those models, the duration of each element is learned via modification of the recurrent connections within each module^11–13^. However, these two components are still not sufficient in order to avoid sequence compression during recall. In order to solve this problem, we assume additional structure within each module and in the allowed connections between modules. This additional structure allows us to avoid compression during recall while using relatively simple biophysically realistic learning rules. The cellular response types generated by this network are consistent with experimental observation^6,7^, as described below.

Our network is composed of different modules that are selectively activated via feed-forward connections by different external stimuli. Within each module there are two populations of excitatory cells, as well as inhibitory cells (Fig. 2a). The excitatory cells in both populations are identical in their intrinsic properties but differ in their learned and fixed connections with other cells within the module and in different modules. We name these two excitatory populations “Timers” and “Messengers” - the reason for these terms will become clear below as we describe their roles within the network. The “Timer” cells have strong recurrent connections with other Timer cells within the module, and they also excite the inhibitory cells (Inh) and the “Messenger” cells. Inhibitory cells in the module decay quicker than their Timer counterparts, thanks to shorter time constants for synaptic activation (80ms for excitatory, 10ms for inhibitory), and small Timer to Inhibitory weights (there are, in general, a number of degenerate sets of parameters which can facilitate quickly decaying Inhibitory cells). The Messenger cells receive excitatory input from Timer cells and inhibitory input from inhibitory cells within the module, and they also learn to project feed-forward connections to Timer cells in other modules.

**Figure 2.**
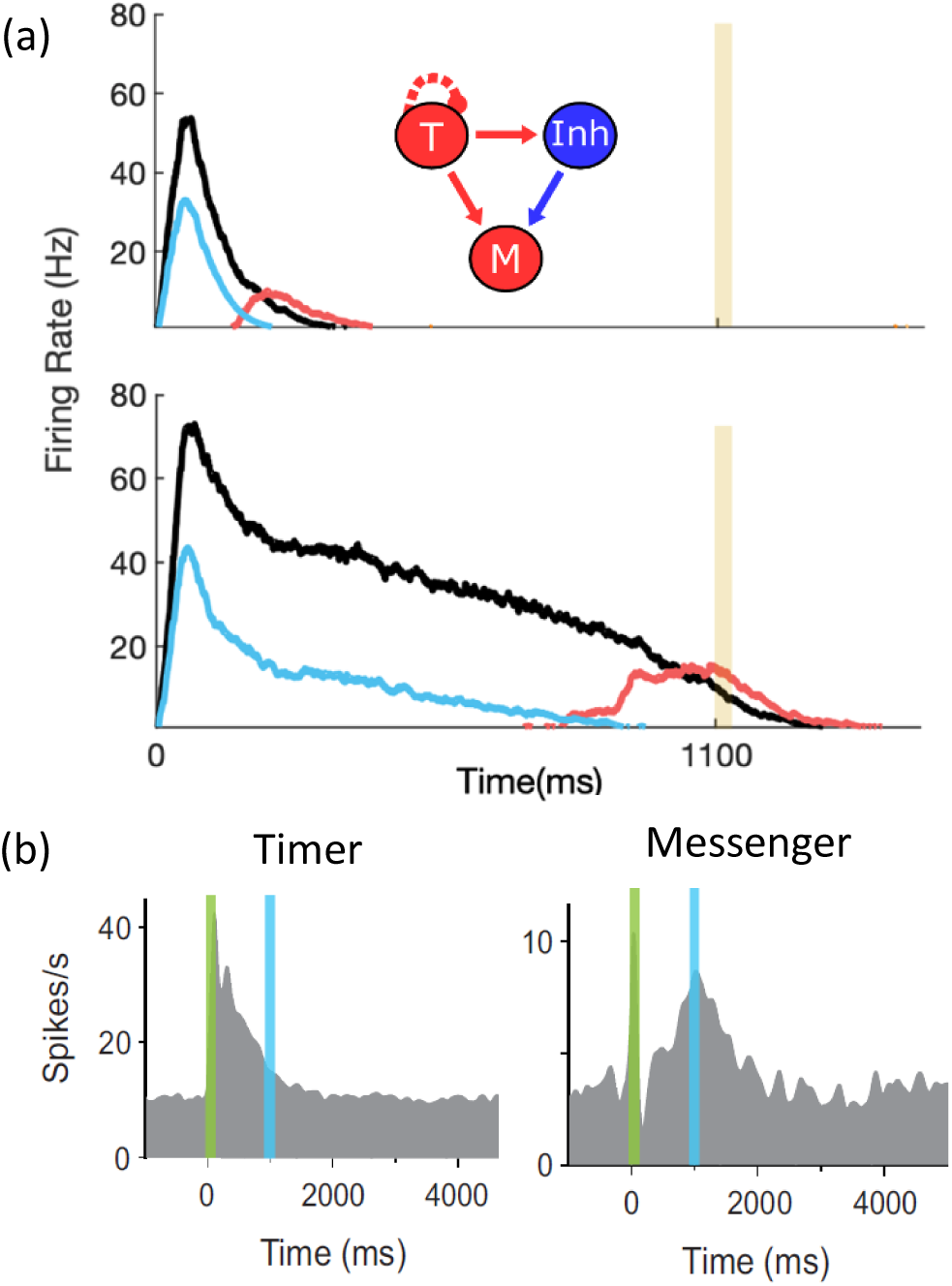
Microcircuit learns time intervals. **a)** Mean firing rates of Timer (black), Messenger (red), and inhibitory populations (light blue) in a microcircuit before learning (top), and after learning (bottom) to represent an 1100 ms interval. Inset: CNA microcircuit. Solid lines indicate fixed connections, while dotted lines indicate learned connections. **b)** Timer and Messenger cell type responses to delayed reward task in V1. Green bar represents stimulus and blue bar represents reward. Figure adapted form Liu et al (2015) with permission.

In this paper we use a previously described reinforcement learning rule based on two competing, Hebbian-activated eligibility traces^24,25^: one for long-term potentiation (LTP) and one for long term depression (LTD). In addition, we have assumed that on every transition between stimuli, a ‘neuromodulator’ is released globally, and this serves as a reinforcement signal (see Methods). We have used this rule because it can solve the temporal credit assignment problem, allowing the network to associate events distal in time, and because it reaches fixed points in both recurrent and feedforward learning tasks^24^. The assumption of a temporally precise but spatially wide spread neuromodulatory signal might seem at odds with common notions of temporally broad neuromodulator release, but they are indeed consistent with recent recordings in several neruomodulatory systems^26,27^.

Using this rule and the described architecture within a single module, we find (Figure 2a) that the network naturally generates cells that learn to be active for the duration of the stimulus (Timers), and other cells that activate towards the end of the stimulus duration (Messengers). The Timer cells learn the duration of the stimulus via changes in the excitatory synaptic efficacies within the Timer population. The Messenger cells evolve their specific temporal profile due to the temporal profiles of their fixed excitatory and inhibitory inputs from the Timer cells and the inhibitory cells. Since the activity of the inhibitory cells decays at a quicker rate than the Timer cells, the Messenger cells arise at the end of the decay phase, when excitation from the Timer cells dominates. These results are consistent with experimental observations in V1 circuits that learn the duration between a stimulus and a reward^6–8^, as shown in Figure 2b. These results replicate our previous results^28^ which used a learning rule with a single trace^29^.

This modular microcircuit, in which both Timer and Messenger cells emerge from learning, acts as a “Core Neural Architecture” (CNA)^28^, an elemental package of basic temporal neural responses which can then be used within the larger network to create more complicated representations. This CNA functions as the basic microcircuit for our sequence model, and each “element” of an input sequence is represented by one of these CNA circuits. One must note that the individual CNAs are plastic, as their temporal properties are not fixed but adapt to the environment. In a structure akin to the microcircuits found in cortical columns^9^, this model places distinct CNAs sensitive to particular visual stimuli in a columnar structure, as shown in Figure 3a. When presenting a sequence of visual stimuli, different CNAs in turn become activated in sequence. Timer cells within those CNAs learn the duration of their particular stimuli via recurrent connections, while order is learned via feed forward connections from Messengers in one column to Timers in a subsequently presented column. We will show that this modular architecture can overcome the problem of sequence compression encountered by “chain-like” models.

**Figure 3.**
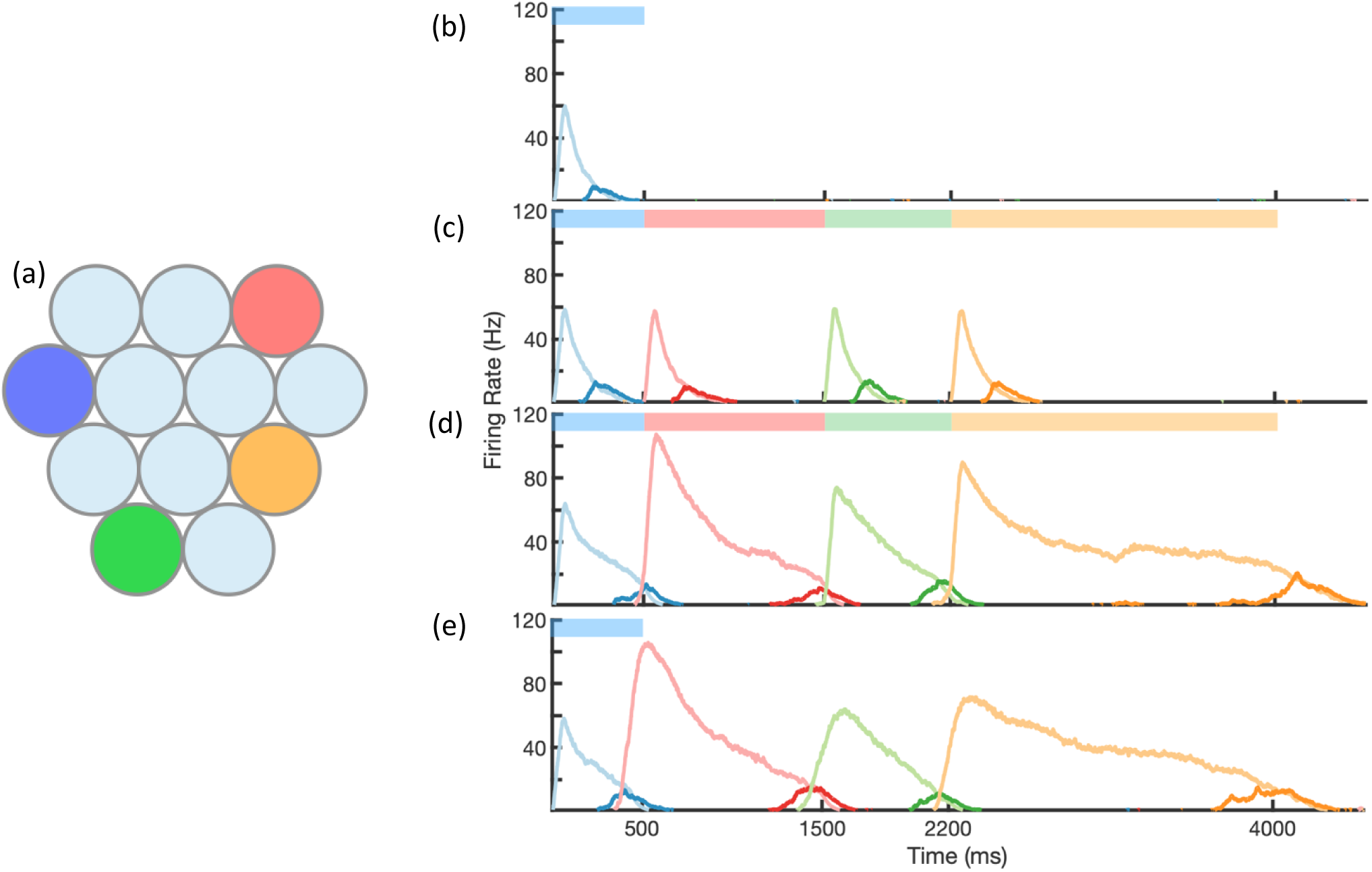
Sequence Learning and Recall. a) Network of ten columns, each containing a CNA microcircuit selective for a different stimulus. Columns containing microcircuits responding to blue, red, green, and orange stimuli are indicated. b) - e) Mean firing rates for Timer cells (light colors) and Messenger cells (dark colors) of 4 different columns during different stages of learning. Stimuli presented are shown as color bars in the top of plots. During learning, columns are stimulated in the sequence indicated by the colorbars (500, 1000, 700, and 1800 ms for blue, red, green, and orange, respectively). b) Before learning, the stimulation of a particular column only causes that column to be transiently active. c) During the first trial of learning, all columns in the sequence become activated by the stimuli, but have not yet learned to represent duration (through recurrent learning of Timer cells) or order (through feed forward learning of the Messenger cells). d) After many trials, the network learns to match the duration and order of presented stimuli. e) After learning, presenting the first element in the sequence is sufficient for recall of the entire sequence.

### Learning and Recalling the Order and Variable Duration of Presented Sequences

With the described architecture and learning rule (see Methods), the network is capable of robustly learning sequences of temporal intervals. Figure 3 shows the network learning a sequence of four different elements, of duration 500ms, 1000ms, 700ms, and 1800ms. The different stimuli here are labeled by color (blue, green, red, orange), and there are four corresponding columns in the network (out of ten) which are sensitive to these stimuli. In this example, the inputs are modeled after LGN responses to spot stimuli^30,31^ (see Supplementary Figure 1), but more generally one may consider these stimuli to be oriented gratings, specific natural images in a movie, or non-visual stimuli such as pitches of sounds in a song.

Before learning, presentation of the blue stimulus only produces a transient response in the CNA microcircuit housed in the blue-sensitive column (Fig. 3b). During learning (Fig. 3c,d), a sequence of stimuli is presented, and microcircuits in their respective columns learn to represent the duration of the stimulus which activates them. The different microcircuits also learn to “chain” together in the order in which they were stimulated. After learning, presentation of the blue stimulus triggers a ‘recall’ of the entire sequence (Fig. 3e): blue for 500ms, red for 1000ms, green for 700ms, and orange for 1800ms. The network is capable of learning sequences of temporal intervals where the individual elements can be anywhere from ∼300ms to ∼1800ms (see Supplementary Figure 2) in duration, which agrees with the observed ranges used for training V1 circuits^1,6^. In simulations we have been able to learn a sequence of up to eight elements (Supplementary Figure 3). The upper limit on the number of elements which can be learned in a sequence has not been fully explored due to the computational time required. For this work, sequences of four elements were chosen to match experimental results^1^. The model presented here is a high dimensional spiking model, but similar results are obtained with a low dimensional rate model in which the activity of each population is presented by a single dynamical variable (for details, see Methods and Supplementary Figure 4).

To gain an understanding of why learning succeeds in this model, we focus on the learning occurring between and within two selected modules in a sequence (Figure 4). Before training, presentation of an isolated stimulus only evokes a transient response of the CNA sensitive to that specific stimulus, since the Timer cells have yet to learn any recurrent connections, and the Messenger cells have yet to establish any feed-forward connections to the Timer cells in other columns (Figure 4a). During training, a particular sequence of inputs is presented, and then repeated over many trials. Hebbian activity in the network triggers activation of synapse-specific LTP and LTD associated eligibility traces, which are then converted into changes in synaptic connections upon neuromodulator release (purple arrows in Figure 4), occurring here closely after a change in stimuli (see Equations 8,9 and 14 in Methods). Each trial pushes the weights in the network towards their fixed points (Equations 17 & 18 in Methods). Hebbian activity within the Timer population causes activation of their respective eligibility traces, and subsequently an increase in their recurrent connections (Supplementary Figure 5). As these lateral connections grow, Timer cells sustain their activity for longer, “extending” their firing profile out in time towards the neuromodulator signal associated with the start of the subsequent stimulus. As this occurs, Messenger cells get “dragged” along by the Timer cells, eventually coactivating with Timer cells of the column which is stimulated next (Figure 4b). This Hebbian coactivation triggers the eligibility traces of these feedforward synaptic connections, before they too are converted into synaptic weight changes by the neuromodulator “novelty” signal (Supplementary Figure 6). After many trials (50-100 for the examples in this paper), weights in the network reach their steady state values (see Methods, Supplementary Figure 2) and learning is complete. As the result of successful learning, a physical synaptic pathway has been traced out which encodes both the duration and order of the input. After learning, the encoded sequence can be recalled by stimulation of only the first element (Figure 4c). Recall does not demonstrate sequence compression, as the Messenger cells’ activation (and thereby the activation of the next element) is appropriately temporally delayed.

**Figure 4.**
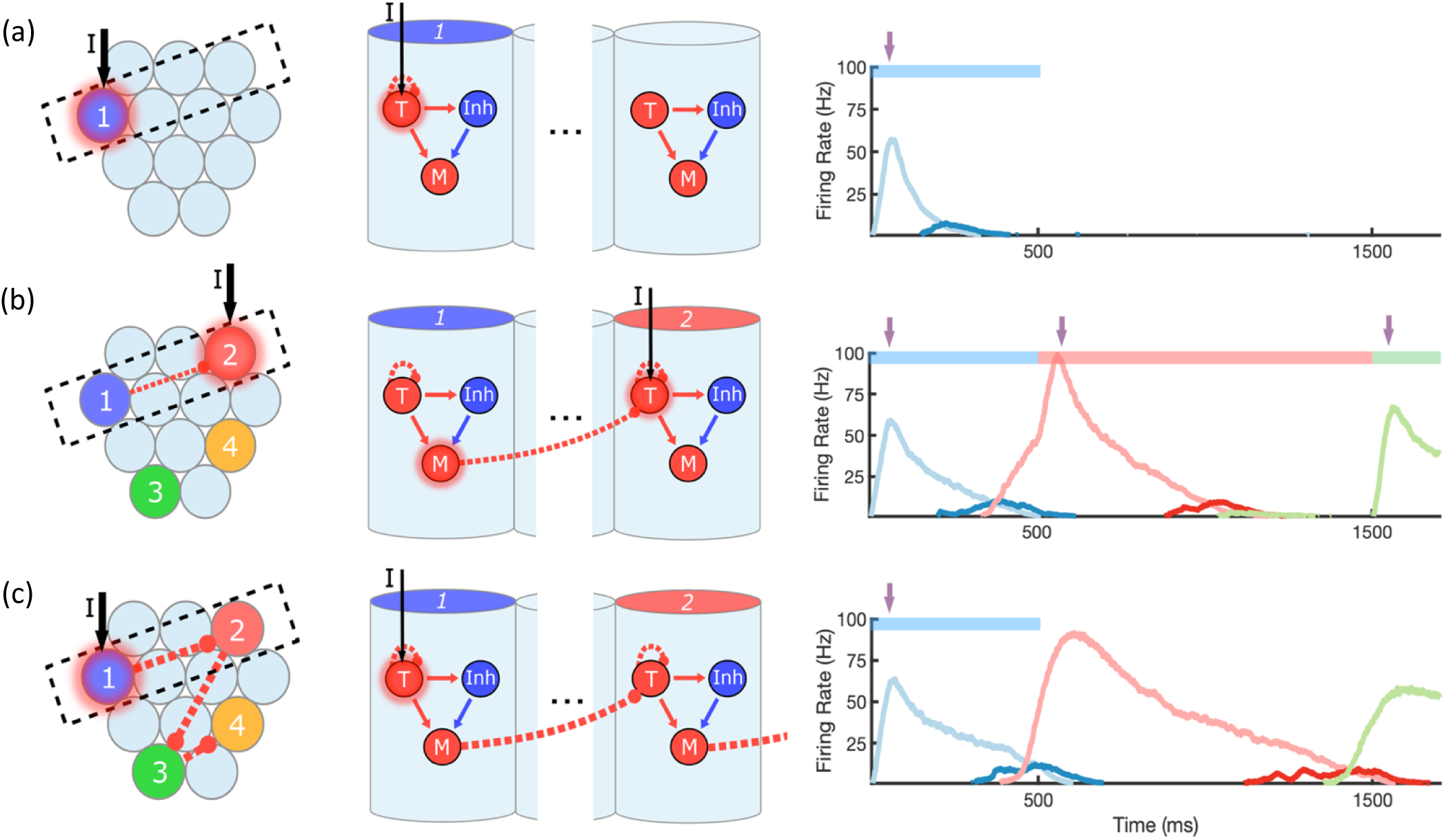
Change in connectivity patters resulting from learning. a) before, b) during, and c) after learning a sequence. Left, view of columnar structure and intercolumnar connections. Dotted box indicates region shown in side view, middle. Middle, the detailed view of two columns and their inter and intracolumnar connectivity. Dotted lines indicate learned connections, continuous lines indicate fixed connections. Right, illustration of mean firing rates for color coded columns. Light colors indicate Timer cells, dark colors indicate Messenger cells. Colorbars indicate stimulated columns. Purple arrows indicate global neuromodulator release. a) Before learning, stimulus of a column’s Timer (T) cells only causes that column to be transiently active. b) If another column is stimulated shortly after the first, the Messenger (M) cells of the previous column will be coactive with the Timer cells of the stimulated column, thereby increasing the feed forward synaptic weights between these two populations. c) After learning, a physical synaptic pathway has been traced out which links columns in the temporal order in which they were stimulated during training.

Properly encoded durations and orders are the result of the fixed points in the learning rule, as described in the Methods section and in previous publications^24,32^. Recurrent learning ends in a fixed point which sets the time D between the end of firing in one column and the start of firing in the next. Feed forward learning results in a fixed point which determines the connection strength between Messenger and Timer cells in subsequent columns. Formally, the fixed point sets the value of the Hebbian term, H_ij_ = *r*_*i*_ · *r*_*j*_, between the Messenger cells and the Timer cells in the next column, and implicitly this results in setting the connection strengths W_ij_. Both these fixed points depend on the parameters of the learning rule, which can be chosen such that these terms achieve desired values (D arbitrarily small, H_ij_ to a fixed value). Such learning can then correctly encode any presented sequence.

Empirically, temporal accuracy of recall depends on many non-trivial factors (i.e. length of individual elements, length of entire sequence, placement of short elements near long elements, etc.), owing to the many non-trivial effects of stochasticity of the spiking network (spike rasters are shown in Supplementary Figure 7). In addition to fluctuations in recall accuracy, there can also be fluctuations in learning accuracy, as randomness in spiking can happen to accumulate such that the traces (and therefore the fixed points) are also sufficiently modified over the course of training. Over the whole network, over the course of many trials, and over the course of learning instances, these effects tend to wash out. Reported times in the recalled sequence generally match the times of the input sequence to within 10% (see Supplementary Figure 2). This model does not observe any intrinsic bias towards over or under-predicting, as this can be modulated by network parameters, and can change stochastically from trial to trial or from element to element.

While the results shown in this work were obtained using a two-trace learning rule (described below), the network can also be trained with a learning rule based on a single trace^29,32^ and the results are similar to those demonstrated here (one-trace results not shown).

### Learning and Recalling non-Markovian sequences

In this modular network, the transitions from each module to the next depend only on the identity of the current module that is active; such a model is formally called a Markovian model^33^. While the Markov property is typically discussed in the context of probabilistic models, here we apply it to deterministic sequences as well. A Markovian model can only reproduce specific types of sequences and is unable to reproduce a sequence in which the same element is repeated more than once and is followed each time by a different element, for example the sequence ABAC. The columnar network described above is essentially Markovian since activation of a neural population will necessarily feed forward to all other populations it is connected to, no matter the history or context. To learn non-Markovian sequences, we modify the network structure. Ideally, the network should be able to learn sequences which are in themselves non-Markovian (ABACAD), as well as simultaneous combinations of sequences which are non-Markovian when learned together (ACE and BCD). We will demonstrate both cases here.

For the network to learn and recall non-Markovian sequences, it must somehow keep track of its history, and use this history to inform transitions during sequence learning and recall. To this end, we include two additional “stages” to the network (Figure 5). The first is a fixed (non-learning) recurrent network, sometimes called a “reservoir” (as in reservoir computing) or a “liquid” (as in a liquid state machine)^34,35^, which receives inputs from the Messenger cells in the main columnar network. Owing to these inputs and due to its strong recurrent connectivity, this network possesses a long-term memory of the state of the columnar network. The second additional stage is a high dimensional, sparse, non-linear network which receives input from the reservoir, serving to project the reservoir states into a space where they are highly separated and non-overlapping. The result is that a given “pattern” in this sparse network at time t uniquely identifies the history of the main network up to and including time t. Since these patterns are highly non-overlapping, a particular pattern at time t can use simple Hebbian learning to connect to Timer cells firing at time t + Δt in the main network (Messenger to Timer feed forward learning is removed in this non-Markovian example). After learning, any time the same particular pattern is recalled (which only occurs if the dynamic history of the network is the same as it was during training), the Timer cells which are thereby stimulated by that pattern will be the same ones which were externally stimulated following that pattern during training.

**Figure 5.**
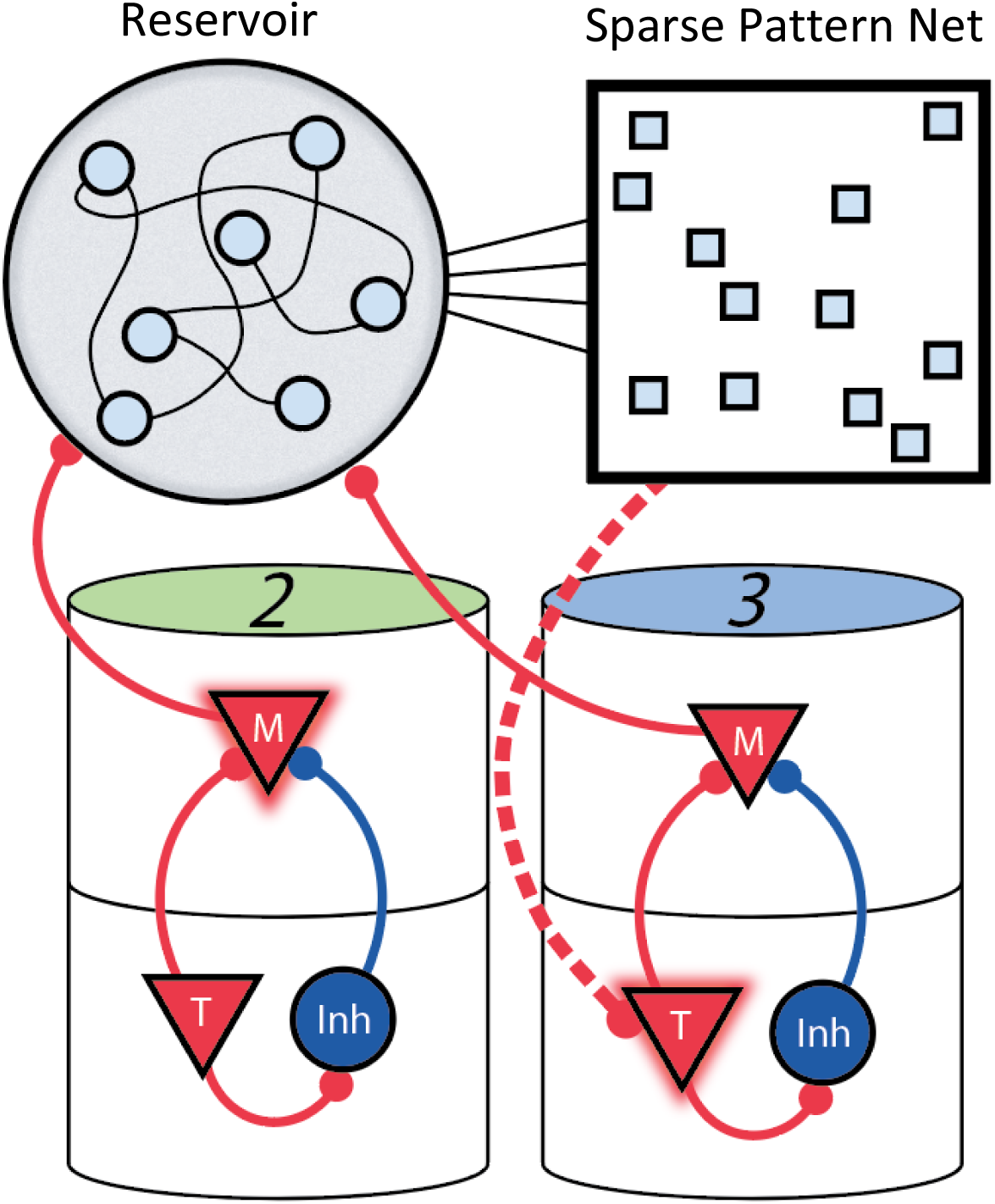
Three-Stage Network. Two sequentially activated columns (2-3) learn to connect to each other through a reservoir and sparse pattern net. At time t, Messenger cells from column 2 are active and act as inputs into the reservoir (earlier, Messenger cells from column 1 also fed into the reservoir). The sparse pattern net receives input from the reservoir, so as to be a unique representation of the history of the network up to and including time t. Timer cells active at t + Δt (column 3) connect to the sparse pattern via Hebbian learning.

We test the ability of this three-stage network to learn long, non-Markovian sequences with repeated elements by presenting a sequence blue-green-blue-red-blue-orange (BGBRBO) during training (Figure 6). A simplified, rate-based version of the columnar network is used to reduce computation time (see Methods). The presented sequence is such that there are three different transitions from blue to another element, and each stimulus is also presented for a different duration, making this a non-trivial problem. Before learning (Figure 6a), the blue stimulus does not evoke any recall, only triggering a transient response of the blue-responsive column. During learning (Figure 6b,c), Timer cells learn the duration of their stimulus, as before, but now it is the sparse pattern net which is using Hebbian learning to feed forward to Timers in subsequently stimulated columns. After learning (Figure 6d), the blue stimulus is sufficient to trigger recall of the entire trained non-Markovian sequence. We also demonstrate an example of simultaneous learning of two sequences with a shared element (Figure 7). First, a sequence blue-red-orange (BRO) is trained, followed by a sequence green-red-purple (GRP). Each of the elements in these sequences has a duration of 500ms. With both these sequences learned and stored simultaneously in the same network, transitions from red are non-Markovian – red should transition to orange if it was preceded by blue, and should transition to purple if it was preceded by green. Figure 7a shows recall upon stimulation of blue, while Figure 7b shows recall upon stimulation of green. In both cases, the sequence makes the correct transition from red to the appropriate third element. Recall of both sequences is also robust to perturbations in the initial state of the reservoir, as shown in Supplementary Figure 8.

**Figure 6.**
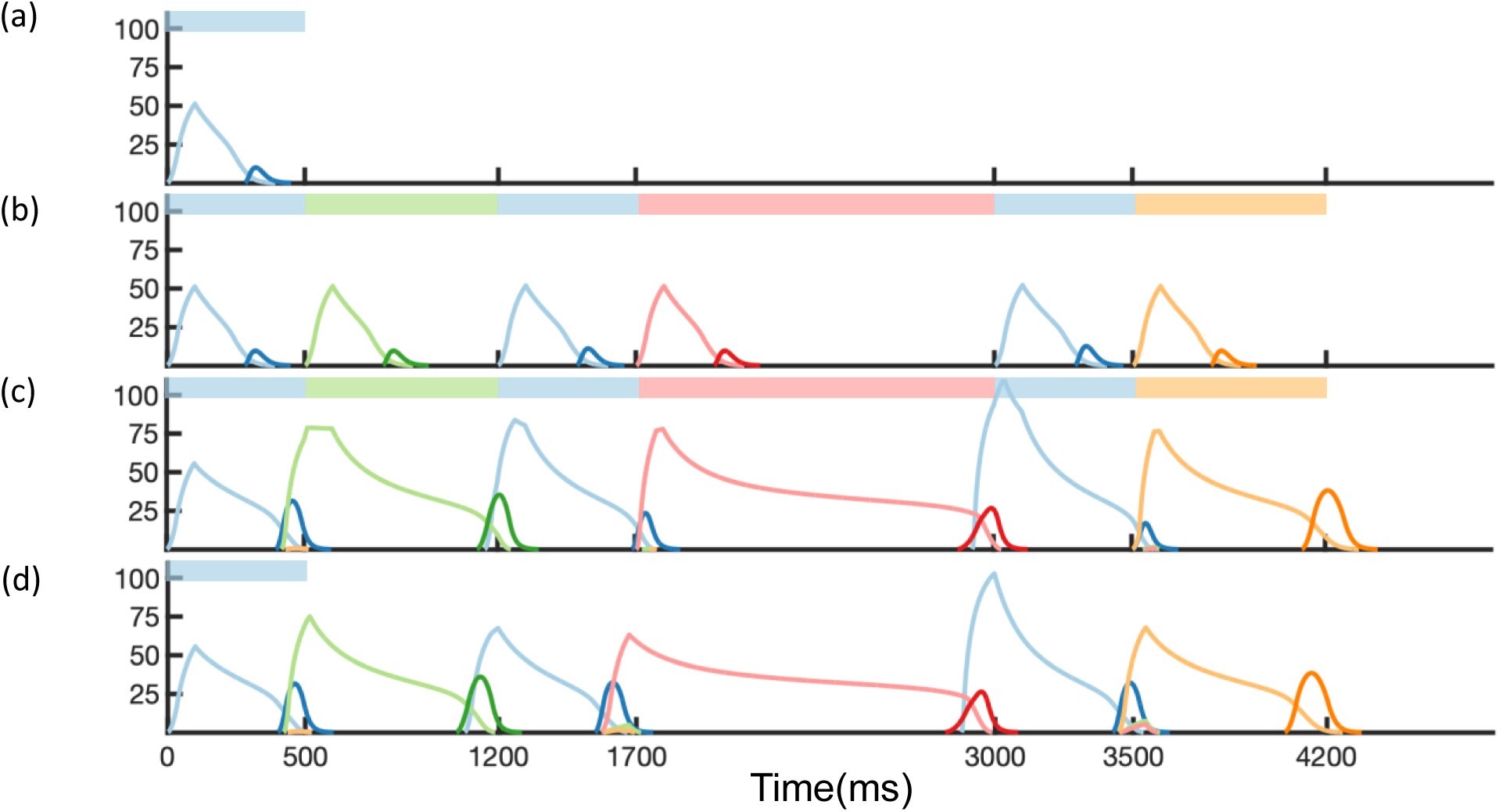
Non-Markovian Sequence Learning and Recall. Mean firing rates for Timer cells (light colors) and Messenger cells (dark colors) of 4 different columns during different stages of learning (before, first trial of learning, last trial of learning, after learning). Stimuli presented are shown in color bars inset in top of plots (500, 700, 500, 1300, 500, and 700 ms for blue, green, blue, red, blue and orange, respectively).

**Figure 7.**
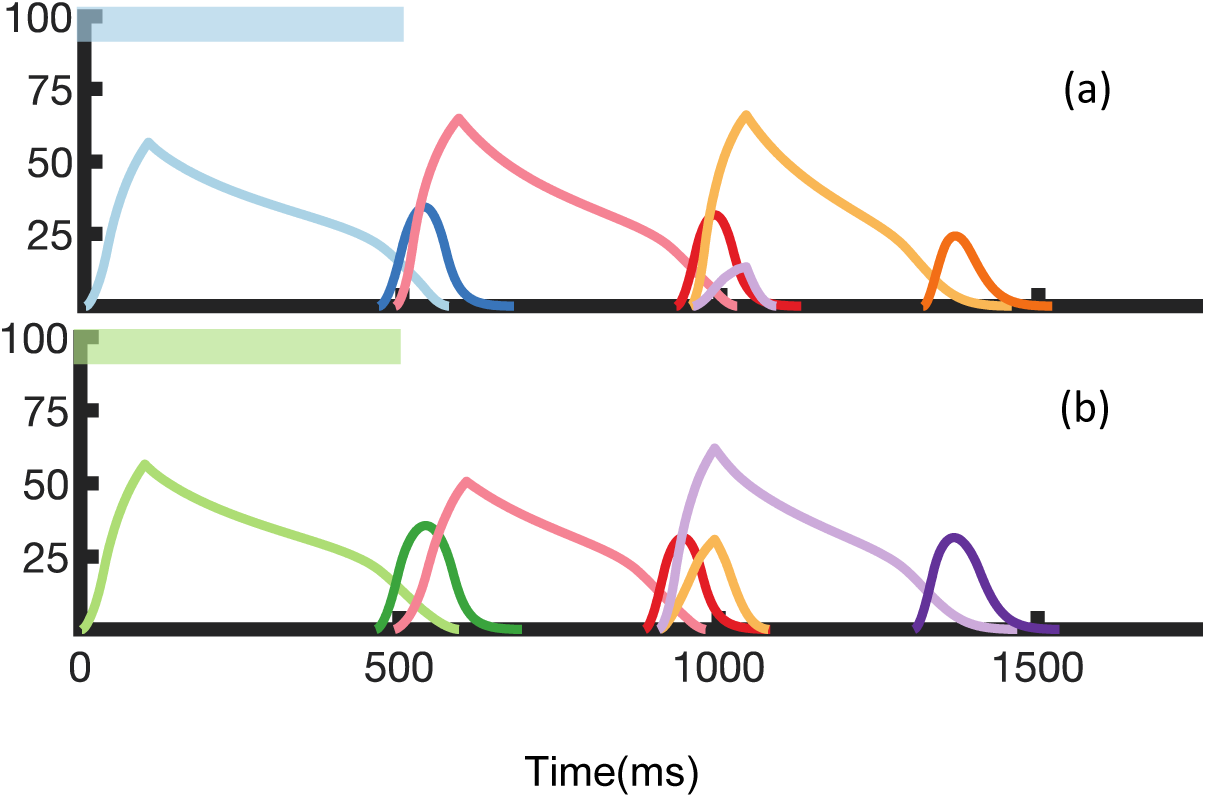
Recall of Two Overlapping Sequences. Mean firing rates for timer cells (light colors) and messenger cells (dark colors) during recall of two sequences. Both blue-red-orange (BRO) and green-red-purple (GRP) have been stored in the network via learning. a) Recall of BRO, following presentation of a blue stimulus. b) Recall of GRP, following presentation of a green stimulus. Note that R transitions to a different element in the two sequences.

Of note in these two simulations is that the “incorrect” transitions do get slightly activated, as the sparse patterns responsible for these transitions are not completely non-overlapping. There is an interplay here between the robustness and uniqueness of these patterns – the sparser they are, the more non-overlapping they are, but also the more sensitive they are to noise. Other parameters in the model, such as the level of recurrence in the reservoir, or the projection strengths from the reservoir to the sparse net, also effect the robustness/uniqueness of the patterns. A full quantification of the robustness, capacity, basin size etc. of the patterns is beyond the scope of this work. For the simulations shown, a set of parameters (Supplementary Table 2) was empirically chosen so that a handful of non-Markovian transitions could be reliably encoded.

The addition of the reservoir and sparse network is essential for the ability to encode and replay non-Markovian sequences, but they alone are not sufficient for sequence learning and recall. One must note that the reservoir receives its input from the modular network, not from the environment, and therefore if the modular network does not learn, the inputs to the reservoir would not be generated appropriately. If instead, we supplied input directly to the reservoir, recall would not be possible because input to the reservoir on learning and recall trials would be very different and therefore the reservoir dynamics would completely differ.

Other groups have approached non-Markovian sequences by including additional hierarchy or long synaptic time constants^36,37^, and solutions like RNNs handle such sequences natively since they are continuous systems with a dependence on a relatively long history when compared to the intrinsic time constants. We combine these two methods, using highly recurrent networks in the context of a larger architecture, and still maintaining biophysical realism by using Hebbian learning rules.

## Methods

### Spiking Dynamics and Two-Trace Learning (TTL) Rule

The network is comprised of microcircuits of both excitatory and inhibitory spiking leaky-integrate-and-fire (LIF) neurons placed in a modular architecture akin to that observed in cortical columns. The following equations describe the membrane dynamics for each model neuron *i*:

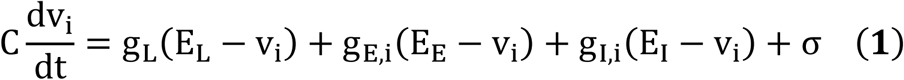

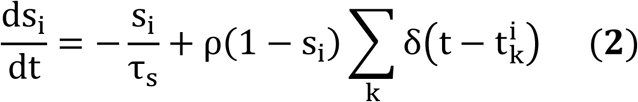

Here, subscripts *L, E*, and *I* refer to leak, excitatory and inhibitory, respectively. *g* refers to the corresponding conductance, and *E* to the corresponding reversal potentials. *v*_*i*_ and *s*_*i*_ are the membrane potential and synaptic activation, respectively, of neuron *i*. σ is a random noise term. Once the membrane potential reaches a threshold *v*_*th*_, the neuron spikes and enters a refractory period *t*_*ref*_. Each spike (at time 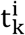) updates the synaptic activation s_i_ by an amount ρ(1 − s_i_), and in the absence of spikes, synaptic activation decays exponentially. Conductance *g* is the product of the synaptic weight matrix with the synaptic activation, summed over all pre-synaptic neurons:

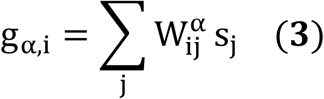

Here, α can be either *E* (excitatory) or *I* (inhibitory), and 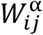 are the connection strengths from neuron *j* to neuron *i*. A firing rate estimate for each neuron r_i_ is calculated as an exponential filter of the spikes at times 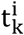, with a time constant τ_r_.

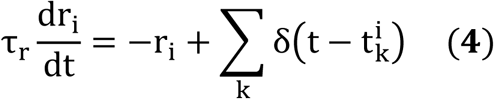

For the rate-based version of these dynamics (used in the three-stage network model), each population of spiking neurons is represented by a single rate-based unit, which is governed by the following equations:

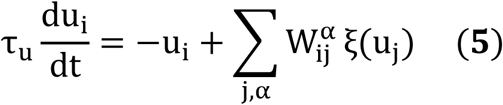

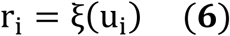

where τ_u_ is the characteristic time constant, and r_i_ is the resulting firing rate for neuron i. ξ is a piecewise nonlinear transfer function^38^:

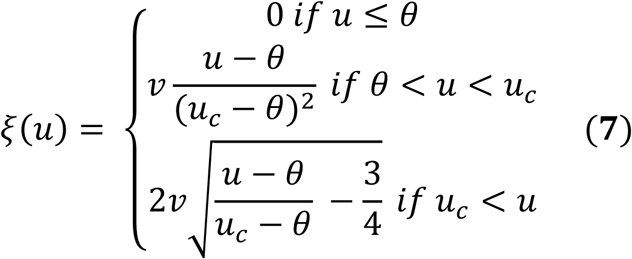

In place of a pair based spike timing dependent rule or rate based Hebbian rule, which fail to solve the temporal credit assignment problem, the network learns based on “eligibility traces” for both long-term potentiation (LTP) and long-term depression (LTD)^28,39^. These eligibility traces are synapse-specific markers that are activated via a Hebbian coincidence of activity between the pre- and post-synaptic cells. At a maximally allowed activation level, these traces saturate, and in the absence of Hebbian activity, these traces decay. LTP and LTD traces are distinct in that they activate, decay, and saturate at different rates/levels. These dynamics are described in the equations below.

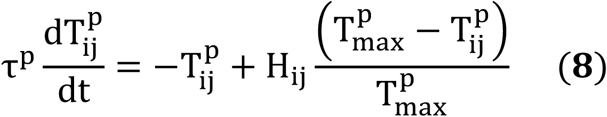

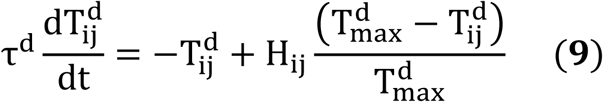

The superscripts p and d indicate LTP or LTD synaptic eligibility traces, respectively. Here, 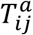 (where *a* ∈ (*p, d*)) is the eligibility trace located at the synapse between the j-th presynaptic cell and the i-th postsynaptic cell. The Hebbian activity, H_ij_, is a simple multiplication *r*_*i*_ · *r*_*j*_ in this rule, where *r*_*j*_ and r_i_ are the time averaged firing rates at the pre- and post-synaptic cells. Here we use a simple case where H_ij_ for LTP and LTD are identical, this is the simplest option and it is sufficient here, but experimentally the “Hebbian” terms are more complex and are different for LTP and LTD traces^25^. The parameter 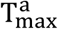 refers to the saturation level, and τ^a^ to the characteristic decay time constant of the trace. If we assume steady state activity such that < *H*_*ij*_ > is constant, we can derive effective saturation levels and effective time constants 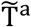 and 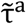, which vary as a function of the Hebbian term:

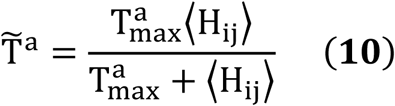

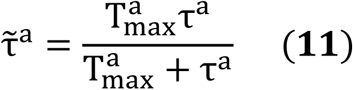

where ⟨H_ij_⟩ is the mean of Hebbian activity during saturation, in cases of prolonged and steady pre- and post-synaptic activation^24^. Equations 8-9, using parameters as in Supplementary Table 1, create traces with a fast, quickly saturating rising phase in the presence of constant activity, and a long, slow falling phase in the absence of activity. The trace dynamics in the rising and falling phases can be approximated using these effective terms to have the simple exponential forms:

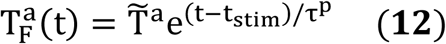

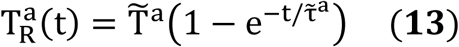

where 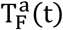 is the trace dynamics in the falling phase, and 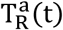 represents the trace dynamics in the rising phase.

In this rule, synaptic weights are updated upon presentation of a global neuromodulatory signal R(t). In general, this signal can be either a traditional “reward” signal, or a “novelty” signal that accompanies the presentation of a new stimulus. This neuromodulatory signal then converts eligibility traces into synaptic efficacies, consuming them in the process, via a simple subtraction of the LTD trace from the LTP trace:

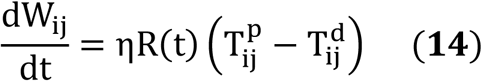

Here, W_ij_ is the synaptic efficacy between pre- and post-synaptic cells j and i, R(t) is the neuromodulatory signal, and η is the learning rate. Following presentation of the neuromodulatory signal, the eligibility traces are “consumed” and reset to zero, and their activation is set into a short refractory period (25ms). Synaptic efficacies reach a steady state under the following condition:

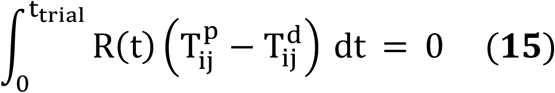

In our model, each time a stimulus starts or ends, a global novelty neuromodulator is released, and this acts as the “reward” in the learning rule. As a result, synaptic efficacies update every time a stimulus is changed, according to the learning rule in Eq. 14, until they reach the fixed point of Eq. 15. Under the simplifying assumption that the reward function R(t) is a delta function δ(t − t_reward_), this fixed point becomes

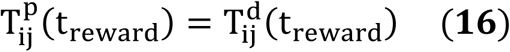

Because of the differing activation, decay, and saturation rates/levels of LTP and LTD traces, this fixed point can be reached either during the rising phase or falling phase (see Supplementary Figures 5 and 6) of the traces. The fixed point of the falling phase corresponds to the recurrent learning of the Timer cells, and the fixed point of the rising phase corresponds to the feed forward learning of the Messenger cells.

To solve for the falling phase fixed point, we first assume that the traces are saturated when the underlying Hebbian activity is above a certain threshold, H_th_. We call the time when Hebbian activity crosses below that threshold t_th_. Then, combining equations 12 and 16 (substituting t_reward_ in for t and t_th_ in for t_stim_), we can solve for D = t_reward_ − t_th_ at the fixed point:

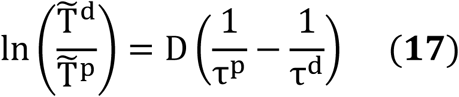

The objective of learning, then, is to move D from its starting value to the value in Eq. 17, which is determined by the parameters in the network. The parameters can be chosen such that the fixed-point value of D is arbitrarily small. As shown in Supplementary Figure 5, this recaptures the behavior of the Timer cells.

The rising phase fixed point can be determined by combining equations 13 and 16 to give us an implicit function of H:

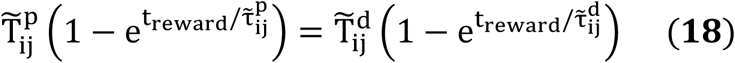

Since 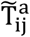 and 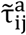 both depend on H (equations 7 and 8), given t_reward_, 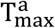, and τ^a^, H is uniquely determined by this fixed point. Practically, this means feed forward learning increases synaptic efficacies W_ij_ in order to increase post-synaptic firing r_i_, until H_ij_ = *r*_*i*_ · *r*_*j*_ = a fixed value, as determined by equation 18 (Supplementary Figure 6).

Through these learning dynamics, the Timer cells learn to represent the duration of their particular element in the sequence (by decreasing D to its fixed-point value), while Messengers learn to feed forward to the Timer cells in the next stimulated column (by increasing W_ij_ until H_ij_ reaches its fixed point value). Note that for modules that do not have overlapping activation, the Hebbian term (H_ij_) is zero, and therefore the associated weights W_ij_ do not change. Supplementary Figure 2c shows these weights approaching their fixed-point values over the course of learning. Earlier work derives these equations in more detail^24^.

The parameters chosen for the traces are displayed in Supplementary Table 1. For the recurrent traces, the parameters follow the restrictions derived from analysis in an earlier publication^24^. The corresponding analysis for the feed-forward traces, however, is not necessarily applicable in the context of sequence learning, since multiple neuromodulator signals are active during each learning epoch. As a result, the parameters for the feed-forward traces are empirically derived. Multiple different sets were found to work, but the set in Supplementary Table 1 was the one used for all simulations included in this paper.

The above eligibility trace learning rules are hereafter referred to as “two-trace” learning or TTL. As described in a previous publication^24^, and demonstrated throughout this work, TTL allows for both feed forward and recurrent learning, and as such can robustly encode temporally dependent input patterns. TTL is supported by recent experiments which have found evidence for eligibility traces existing in cortex^25^, striatum^40^, and hippocampus^41,42^. Other eligibility trace rules, such as the one trace rule demonstrated in earlier work^29,32^, can also replicate the results of this paper. In general, any rule which can associate distal events/rewards, thereby solving the temporal credit assignment problem, would be likely to work with this network model. TTL was chosen for its biological realism, but the novel capabilities of this model (its ability to learn and recall both the duration and order of elements in a sequence) primarily result from the network architecture (described below), combined with a history dependent learning rule.

The three-stage network used to learn and recall non-Markovian sequences is comprised of the main columnar network, a highly recurrent network (reservoir), and a sparse pattern net. The dynamics of the rate-based columnar network are described above. The dynamics of the units u_i_ in the reservoir are described by the equation below:

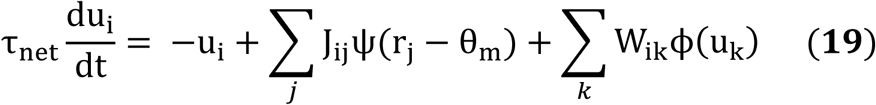

Where u_i_ are the firing rates of the units in the reservoir and r_j_ are the firing rates of the Messenger cells in the main network. J_ij_is a K x n binary matrix of projections from the columnar network to the reservoir, where K is the number of units in the reservoir, and n is the number of columns in the network. J_ij_ is structured such that the first K/n units in the reservoir receive direct input from the Messengers in the first column, the second K/n units receive direct input from the Messengers in the second column, and so on. W_ik_ are the recurrent weights of the reservoir, each of which is drawn from a normal distribution 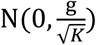, where g is the “gain” of the network^43^. ψ is a piecewise linear activation function:

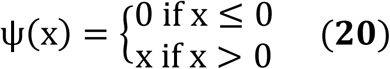

ϕ is a sigmoidal activation function:

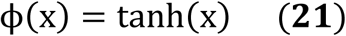

The reservoir projects to a high-dimensional sparse pattern network. Each unit in the sparse pattern net receives input from the reservoir, fed through a sparse matrix O_mi_, with fraction *ρ* = .04 of entries non-zero and drawn from a normal distribution 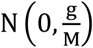, where M is the number of units in the sparse pattern net and g is the gain, as described above. The activation of each unit *i*_m_ of the sparse network is determined by the following:

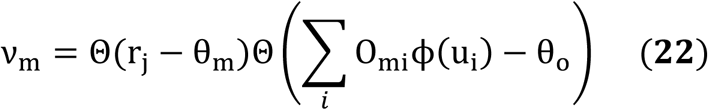

where Θ is the Heaviside function. The sparse pattern network is binary in our implementation but this is not essential, as long as its non-linear and sparse. The sparse network feeds back into the main network via weights Q_jm_, which are learned via a simple Hebbian rule:

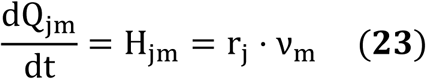

with the additional restriction 0 ≤ Q_jm_ ≤ Q_max_. Here, *j* indexes units in the main network, and *m* indexes units in the sparse pattern net.

### Network Architecture

In a totally unstructured network, the above learning rules would be insufficient for the task of sequence learning and memory. In exchange for the freedom of online, biophysically realistic reinforcement learning, we must presuppose some restrictions (still biophysically realistic) on macro structure in the network. The structure imposed on the network has two components: a modular columnar structure with restricted connections, and the weight distribution of the non-plastic synaptic efficacies. The weight distribution results in the emergence of distinct Timer and Messenger cell types, and the columnar structure places populations of these cells into stimulus-specific modules.

The network consists of ten different “columns”, each containing 100 excitatory Timer cells, 100 inhibitory Timer cells, 100 excitatory Messenger cells, and 100 inhibitory Messenger cells, all assembled in a CNA microcircuit. Each column is tuned to be selective for a particular stimulus from the input layer, such that a sequence of external stimuli (e.g. ABCD or “red green orange blue”) will trigger a corresponding sequence of input-selective columns.

To simulate presentation of visual cues, the network is stimulated by an input layer designed to mimic spiking inputs from the lateral geniculate nucleus. Each stimulus is represented in the input layer by a Poisson spike train with a 50ms pulsed peak, in accordance with the observed cellular responses in LGN^30,31^. In general, this spike train can also include a decaying exponential tail, but short pulses were chosen both for analytical simplicity and to match experimental data^30,31^, as shown in Supplementary Figure 1. The cells of the input layer feed directly into the respective Timer cells of the columnar network.

Intercolumnar inhibition exists between Timer populations and between Messenger populations in the different columns, for the purposes of soft winner-take-all (WTA) dynamics between the columns of the network (Figure 8a). Such inhibition is generally not necessary in the low-noise case (or in a rate-based model), but the stochastic, spiking nature of the network makes it a desirable practical inclusion to guard against spurious runaway excitatory chains.

**Figure 8.**
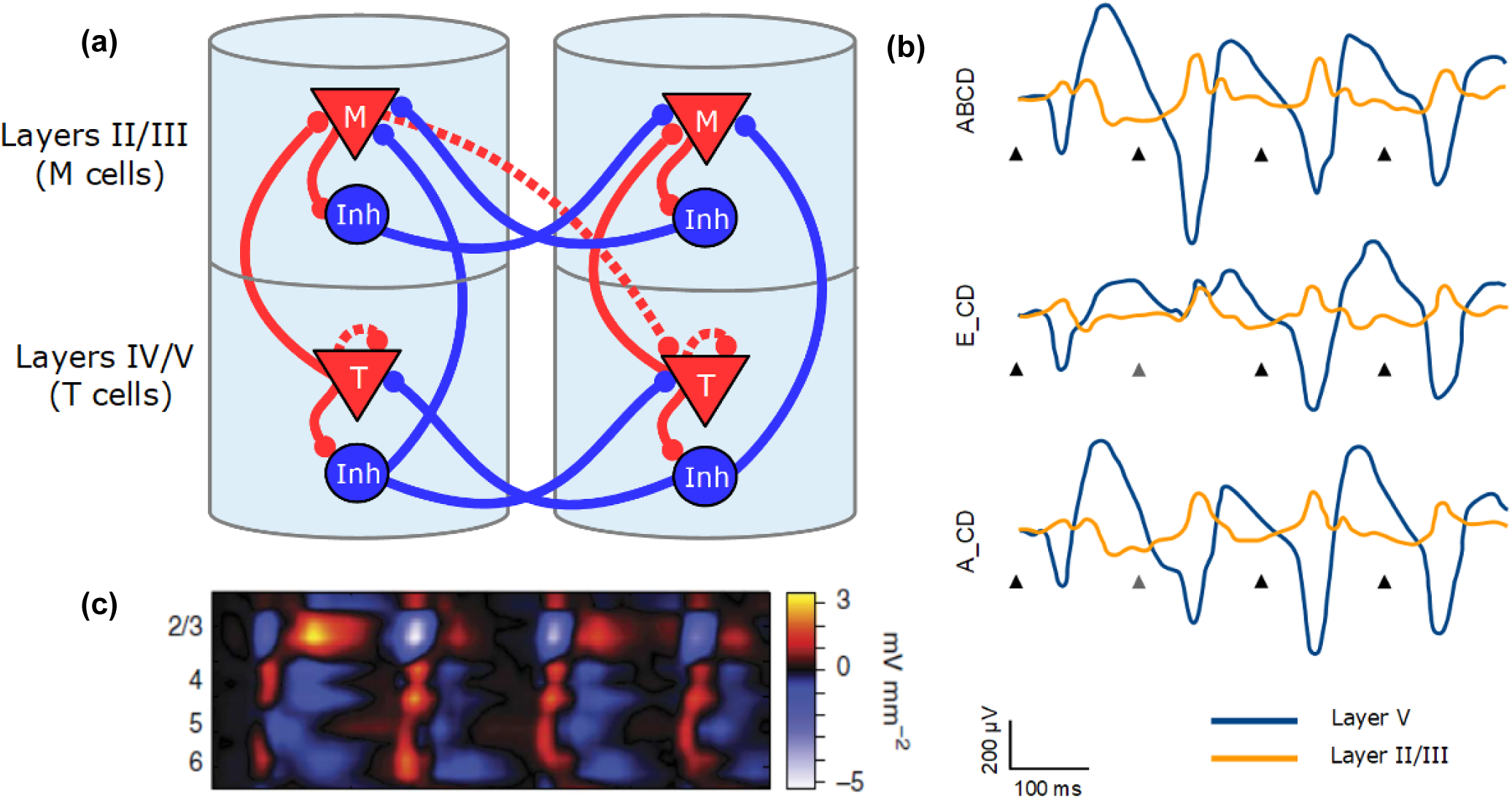
Explicit microcircuit structure. **a)** Complete microcircuit structure displayed in laminar architecture of cortical columns. Dashed lines represented learned connections, while continuous lines represent fixed connections. Intercolumn inhibition produces winner-take all dynamics between columns. **b)** Local field potentials from visual cortex in mice repeatedly exposed to a sequence of oriented gratings, ABCD. Presenting the trained sequence with the second element omitted (A_CD) still appears sufficient for recall of the ABCD network response. Figure adapted from Gavornik and Bear (2014) with permission. **c)** CSD analysis from Gavornik and Bear (2014), showing laminar specificity of responses to sequences ABCD (top), and A_CD (bottom). Note the presence of peaked messenger-like responses in layer II/III and the presence of extended timer-like responses in layers IV/V/VI.

For the purposes of this paper, Timer cells can only learn to connect to other Timer cells within their column, and Messenger cells can only learn connect to (any) Timer cells outside of their column. (Figure 8a). In case of the extended network that can learn non-Markov sequences, Messenger cells connect to the reservoir, and not directly to Timer cells in other columns, as described above. While we include these restrictions for practical purposes, previous publications have shown all possible intracolumnar excitatory connections to be learnable with use of a single trace eligibility rule^28^. The strengths of non-plastic connections within columns are modeled after those learned in previous publications^28^. These fixed connections serve to establish the Timer and Messenger cells used in the network. By only learning task relevant connections, we simplify the computation and analysis while maintaining the complete general functionality of variable sequence learning and recall.

## Discussion

In this work, we demonstrate the ability of a biophysically realistic network to learn and recall sequences, correctly reporting the duration and order of the individual elements. In combining modular, heterogenous structure with a learning rule based on eligibility traces, the model can accurately learn and recall sequences of up to at least 8 elements, with each element anywhere from ∼300ms to ∼1800ms in duration. We have also shown a modified architecture that is capable of learning and recalling non-Markovian sequences with multiple history-dependent transitions.

The capabilities, construction, and rules of our model stand in contrast to many contemporary and historical models of sequence learning, such as synfire chains and RNNs. In particular, we are not aware of another model that uses biophysically realistic spiking networks and learning rules, along with experimentally observed cell responses, to robustly learn both the order and duration of presented sequences of stimuli. Previous approaches which have attempted to learn both duration and order of elements^44^, have been highly sensitive to changes in parameters, and were not based on experimentally observed cell responses. Other types of hybrid models combine chain structures with additional hierarchy/functionality^14,15^, but either require manually set adaptation time constants that are specific for a set duration to represent the duration of elements, or do not treat the duration of elements as variable at all. Recently another model has been proposed, based on a periodic and essentially deterministic timing network, which also uses biologically plausible learning rules to learn to represent single non-Markovian sequences^45^. However, it cannot learn arbitrarily long (and self-terminating) sequences, such as those presented in Figure 6, nor can it learn several different sequences and replay them separately, such as those presented in Figure 7.

The ability of our model to accurately learn and recall sequences is only possible because of the functionally heterogeneous cell types in the CNA microcircuit being given different roles in learning: it is the job of the Timer cells to encode the duration of a particular element, and the job of the Messenger cells (which only fire at the end of the duration), to pass along this information and encode the order in which the elements are presented. Even though this work also provides further validation of Two-Trace Learning, TTL itself is not required, since model also retains its functionality when learning with an eligibility trace rule with only one trace^29^.

The modified three-stage model is capable of learning non-Markovian sequences because of inclusion of a non-learning recurrent network that stores memory over longer durations than the modular network. However, the model’s ability to generate learned sequences depends crucially on the backbone of the modular network – because of this, we can use a non-plastic, highly recurrent reservoir and maintain biophysically realistic learning rules in the network. This modified model also differs in functionality from traditional RNNs, as in our model, the stimulus presented during training (the entire sequence) is different from the stimulus presented after learning in order to trigger recall (usually the first element is sufficient). In general, the number of elements necessary to trigger recall is the same as the number of elements needed to disambiguate the sequence (i.e. “A” would be sufficient to trigger ABCD in a network embedded with learned sequences ABCD and EFGH, but “AB” would be required to trigger ABCD in a network embedded with learned sequences ABCD and AFGH). In a way, our network acts as an autoencoder, both learning a reduced encoding (e.g. “A” instead of “ABCD”), and reconstructing the original external input from this reduced encoding.

The reservoir and sparse network components of our three-stage model could be thought to arise from a projection from other cortical or subcortical areas. Functionally similar networks (ones that take complex, multi-modal, and dynamic context and repackage it into sparse, separated patterns) have been observed in the dentate gyrus^46,47^ and the cerebellum^48,49^. However, these model components could also be thought of as part of the same cortical network, partially segregated in function but not necessarily by location. Pattern separation is likely a common neural computation that might occur in many brain areas, so we make no particular predictions about the locations of these network components.

Our model suggests that the types of single cell responses in V1 observed in delayed reward tasks (Timers and Messengers)^6–8^ will also be present when learning a visual sequence. Furthermore, we predict that distinct populations of these cells are sensitive to and can learn the dynamics of distinct stimuli. This functional modularity could be physically implemented as physically compartmentalized in columns and layers (Figure 8a), though this is not strictly required. Recent results showing sequence learning and recall in visual cortex support the idea that these cell types are implicated in sequence learning^1^. When multiple visual stimuli are presented sequentially, LFP recordings indicate Timer like responses (long, sustained potentiation) in layer IV/V, with Messenger like responses (short bursts centered around the transition between elements) arising in layer II/III (Figure 8b,c). These results are consistent with the hypothesis that Timers and Messengers are indeed compartmentalized into different cortical layers.

Our model makes several testable predictions. It predicts that: 1. After learning a sequence, Messenger cells will more strongly connect to Timer cells that represent subsequent stimuli, than to Timers which represent previous stimuli, or to other Messengers. 2. Sequence learning will increase lateral connections efficacies between Timers within the same module. 3. There will be a population of inhibitory cells within each module that have firing properties similar to Timers but that decay more quickly (Fig 2a).

## Competing interests

The authors declare no competing interests.

## Notes

### Competing Interest Statement

The authors have declared no competing interest.

## References

1. Gavornik, J. P. & Bear, M. F. Learned spatiotemporal sequence recognition and prediction in primary visual cortex. Nat. Neurosci. 17, 732–737 (2014).

2. Xu, S., Jiang, W., Poo, M. & Dan, Y. Activity recall in a visual cortical ensemble. Nat. Neurosci. 15, 449–455 (2012).

3. Cooke, S. F., Komorowski, R. W., Kaplan, E. S., Gavornik, J. P. & Bear, M. F. Visual recognition memory, manifested as long-term habituation, requires synaptic plasticity in V1. Nat. Neurosci. 18, 262–271 (2015).

4. Eagleman, S. L. & Dragoi, V. Image sequence reactivation in awake V4 networks. Proc. Natl. Acad. Sci. 109, 19450–19455 (2012).

5. Yin, P., Mishkin, M., Sutter, M. & Fritz, J. B. Early Stages of Melody Processing: Stimulus-Sequence and Task-Dependent Neuronal Activity in Monkey Auditory Cortical Fields A1 and R. J. Neurophysiol. 100, 3009–3029 (2008).

6. Shuler, M. G. H. & Bear, M. F. Reward Timing in Primary Visual Cortex. Science (2006).

7. Liu, C.-H., Coleman, J. E., Davoudi, H., Zhang, K. & Hussain Shuler, M. G. Selective Activation of a Putative Reinforcement Signal Conditions Cued Interval Timing in Primary Visual Cortex. Curr. Biol. 25, 1551–1561 (2015).

8. Chubykin, A. A., Roach, E. B., Bear, M. F. & Shuler, M. G. H. A Cholinergic Mechanism for Reward Timing within Primary Visual Cortex. Neuron 77, 723–735 (2013).

9. Potjans, T. C. & Diesmann, M. The Cell-Type Specific Cortical Microcircuit: Relating Structure and Activity in a Full-Scale Spiking Network Model. Cereb. Cortex 24, 785–806 (2014).

10. Binzegger, T., Douglas, R. J. & Martin, K. A. C. Topology and dynamics of the canonical circuit of cat V1. Neural Netw. 22, 1071–1078 (2009).

11. Fiete, I. R., Senn, W., Wang, C. Z. H. & Hahnloser, R. H. R. Spike-Time-Dependent Plasticity and Heterosynaptic Competition Organize Networks to Produce Long Scale-Free Sequences of Neural Activity. Neuron 65, 563–576 (2010).

12. Pereira, U. & Brunel, N. Unsupervised Learning of Persistent and Sequential Activity. Front. Comput. Neurosci. 13, (2020).

13. Klos, C., Miner, D. & Triesch, J. Bridging structure and function: A model of sequence learning and prediction in primary visual cortex. PLOS Comput. Biol. 14, e1006187 (2018).

14. Murray, J. M. & Escola, G. S. Learning multiple variable-speed sequences in striatum via cortical tutoring. eLife 6, e26084 (2017).

15. Martinez, R. H., Lansner, A. & Herman, P. Probabilistic associative learning suffices for learning the temporal structure of multiple sequences. PLOS ONE 14, e0220161 (2019).

16. Rajan, K., Harvey, C. D. & Tank, D. W. Recurrent Network Models of Sequence Generation and Memory. Neuron 90, 128–142 (2016).

17. DePasquale, B., Cueva, C. J., Rajan, K., Escola, G. S. & Abbott, L. F. full-FORCE: A target-based method for training recurrent networks. PLOS ONE 13, e0191527 (2018).

18. Laje, R. & Buonomano, D. V. Robust timing and motor patterns by taming chaos in recurrent neural networks. Nat. Neurosci. 16, 925–933 (2013).

19. Sussillo, D. & Abbott, L. F. Generating Coherent Patterns of Activity from Chaotic Neural Networks. Neuron 63, 544–557 (2009).

20. Nicola, W. & Clopath, C. Supervised learning in spiking neural networks with FORCE training. Nat. Commun. 8, 2208 (2017).

21. Whittington, J. C. R. & Bogacz, R. Theories of Error Back-Propagation in the Brain. Trends Cogn. Sci. 23, 235–250 (2019).

22. Lillicrap, T. P., Cownden, D., Tweed, D. B. & Akerman, C. J. Random synaptic feedback weights support error backpropagation for deep learning. Nat. Commun. 7, 13276 (2016).

23. Murray, J. M. Local online learning in recurrent networks with random feedback. eLife 8, e43299 (2019).

24. Huertas, M. A., Schwettmann, S. E. & Shouval, H. Z. The Role of Multiple Neuromodulators in Reinforcement Learning That Is Based on Competition between Eligibility Traces. Front. Synaptic Neurosci. 8, (2016).

25. He, K. et al. Distinct Eligibility Traces for LTP and LTD in Cortical Synapses. Neuron 88, 528–538 (2015).

26. Howe, M. W. & Dombeck, D. A. Rapid signalling in distinct dopaminergic axons during locomotion and reward. Nature 535, 505 (2016).

27. Hangya, B., Ranade, S. P., Lorenc, M. & Kepecs, A. Central cholinergic neurons are rapidly recruited by reinforcement feedback. Cell 162, 1155–1168 (2015).

28. Huertas, M. A., Hussain Shuler, M. G. & Shouval, H. Z. A Simple Network Architecture Accounts for Diverse Reward Time Responses in Primary Visual Cortex. J. Neurosci. 35, 12659–12672 (2015).

29. Gavornik, J. P., Shuler, M. G. H., Loewenstein, Y., Bear, M. F. & Shouval, H. Z. Learning reward timing in cortex through reward dependent expression of synaptic plasticity. Proc. Natl. Acad. Sci. 106, 6826–6831 (2009).

30. Ruksenas, O., Bulatov, A. & Heggelund, P. Dynamics of Spatial Resolution of Single Units in the Lateral Geniculate Nucleus of Cat During Brief Visual Stimulation. J. Neurophysiol. 97, 1445–1456 (2007).

31. Mastronarde, D. N. Two classes of single-input X-cells in cat lateral geniculate nucleus. II. Retinal inputs and the generation of receptive-field properties. J. Neurophysiol. 57, 381–413 (1987).

32. Gavornik, J. P. & Shouval, H. Z. A network of spiking neurons that can represent interval timing: mean field analysis. J. Comput. Neurosci. 30, 501–513 (2011).

33. Gillespie, D. T. Markov Processes: An Introduction for Physical Scientists. (Elsevier, 1991).

34. Maass, W., Natschläger, T. & Markram, H. Real-Time Computing Without Stable States: A New Framework for Neural Computation Based on Perturbations. Neural Comput. 14, 2531–2560 (2002).

35. Maass, W., Natschläger, T. & Markram, H. Fading memory and kernel properties of generic cortical microcircuit models. J. Physiol.-Paris 98, 315–330 (2004).

36. Hawkins, J. & Ahmad, S. Why Neurons Have Thousands of Synapses, a Theory of Sequence Memory in Neocortex. Front. Neural Circuits 10, (2016).

37. Tully, P. J., Lindén, H., Hennig, M. H. & Lansner, A. Spike-Based Bayesian-Hebbian Learning of Temporal Sequences. PLOS Comput. Biol. 12, e1004954 (2016).

38. Brunel, N. Dynamics and Plasticity of Stimulus-selective Persistent Activity in Cortical Network Models. Cereb. Cortex 13, 1151–1161 (2003).

39. Frémaux, N. & Gerstner, W. Neuromodulated Spike-Timing-Dependent Plasticity, and Theory of Three-Factor Learning Rules. Front. Neural Circuits 9, (2016).

40. Yagishita, S. et al. A critical time window for dopamine actions on the structural plasticity of dendritic spines. Science 345, 1616–1620 (2014).

41. Bittner, K. C., Milstein, A. D., Grienberger, C., Romani, S. & Magee, J. C. Behavioral time scale synaptic plasticity underlies CA1 place fields. Science 357, 1033–1036 (2017).

42. Brzosko, Z., Zannone, S., Schultz, W., Clopath, C. & Paulsen, O. Sequential neuromodulation of Hebbian plasticity offers mechanism for effective reward-based navigation. eLife 6, e27756 (2017).

43. Rajan, K., Abbott, L. F. & Sompolinsky, H. Stimulus-Dependent Suppression of Chaos in Recurrent Neural Networks. Phys. Rev. E 82, 011903 (2010).

44. Veliz-Cuba, A., Shouval, H., Josic, K. & Kilpatrick, Z. P. Networks that learn the precise timing of event sequences. 14121713 Q-Bio (2014).

45. Maes, A., Barahona, M. & Clopath, C. Learning spatiotemporal signals using a recurrent spiking network that discretizes time. PLOS Comput. Biol. 16, e1007606 (2020).

46. Leutgeb, J. K., Leutgeb, S., Moser, M.-B. & Moser, E. I. Pattern Separation in the Dentate Gyrus and CA3 of the Hippocampus. Science 315, 961–966 (2007).

47. van Dijk, M. T. & Fenton, A. A. On How the Dentate Gyrus Contributes to Memory Discrimination. Neuron 98, 832-845.e5 (2018).

48. Chadderton, P., Margrie, T. W. & Häusser, M. Integration of quanta in cerebellar granule cells during sensory processing. Nature 428, 856–860 (2004).

49. Billings, G., Piasini, E., Lőrincz, A., Nusser, Z. & Silver, R. A. Network Structure within the Cerebellar Input Layer Enables Lossless Sparse Encoding. Neuron 83, 960–974 (2014).

