## Supplemental Figures for "Learning precise spatiotemporal sequences via biophysically realistic circuits with modular structure"

### Supplementary Figures

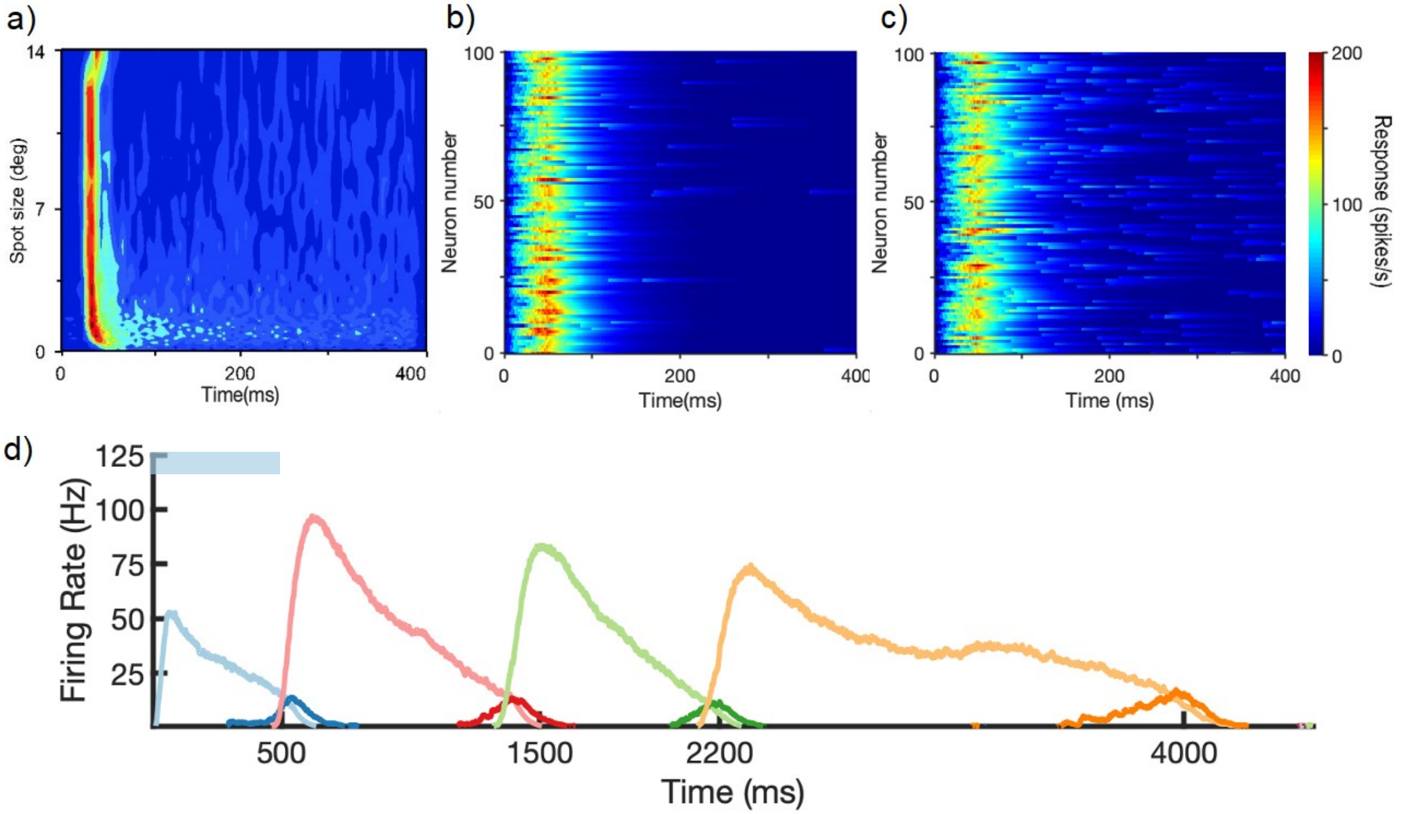


**Supplementary Figure 1. Input Layer Dynamics.** a) Extracellular recording from Ruksenas et al (2007)[30] showing the response of an LGN neuron to a 400ms stimulus, over differing spot sizes. Notably, this LGN neuron only transiently fires, even to a stimulus presented for an extended time. b) Simulations showing response of input layer units to 400ms stimulus (fixed spot size, 7 degrees). The input is approximated as a 50ms pulse of Poisson spikes. This is the approximation used in the main figures of the paper. c) Same as b), but with input approximated by a 50ms pulse of Poisson spikes followed by a decaying exponential tail of Poisson spikes for the remainder of the stimulus time. d) Recall of a learned sequence, in a network with the “pulse+decaying tail” input structure, as in c).

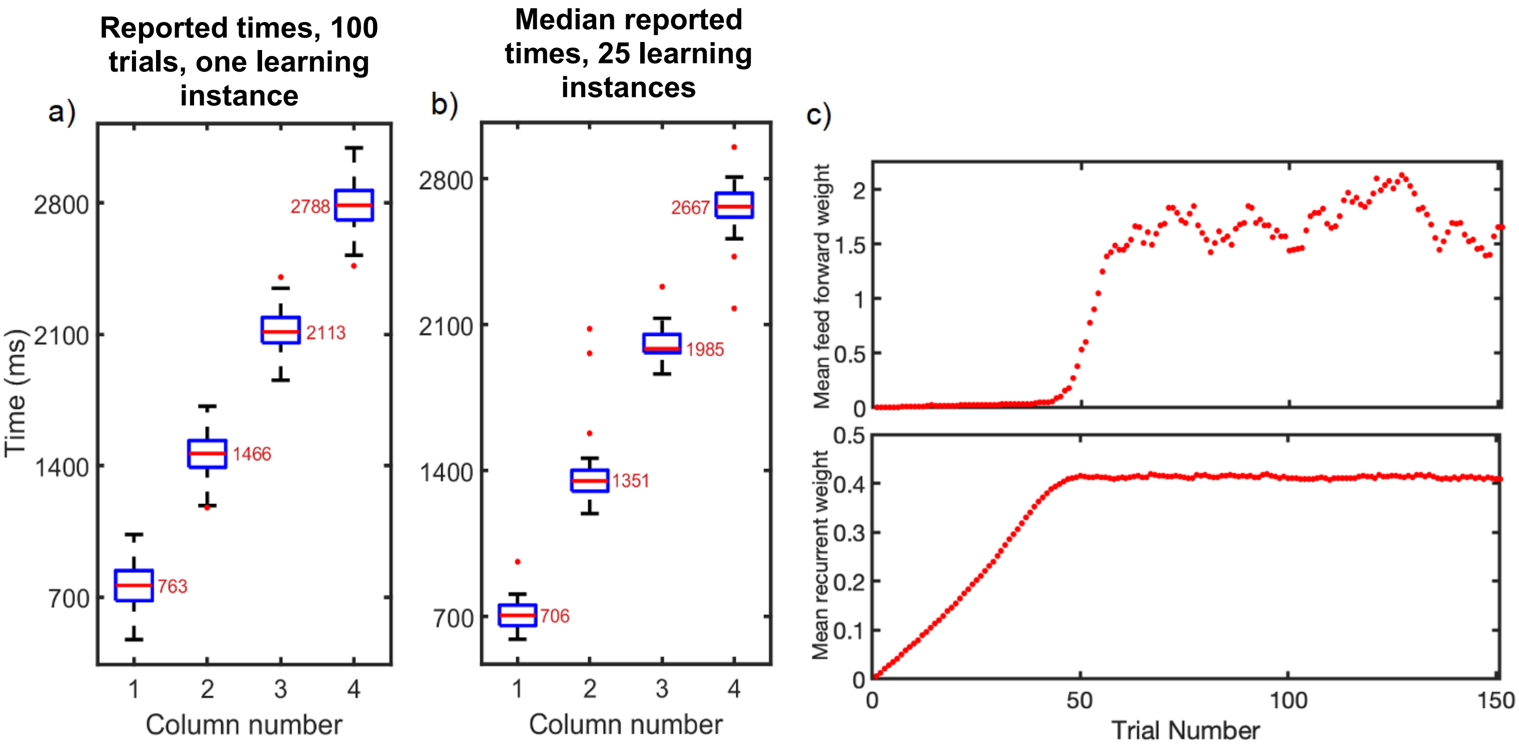


**Supplementary Figure 2. Accuracy of Learning and Recall.** A network is trained to a sequence of 4 elements, each 700ms in duration. Owing to stochastic nature of spiking network, reported times can fluctuate from presentation to presentation, and from learning instance to learning instance. “Reported time” is the time at which the sequence, up to and including that column, drops below 10 Hz. **a)** Recall fluctuations. Reported times for 100 trials of recall, after one learning instance. Median reported time indicated by red bar. Top and bottom of box indicate 25$^{th}$ and 75$^{th}$ percentiles of reported times. The whiskers indicate the maximum and minimum reported times not considered outliers. Red dots indicate outliers. **b)** Learning fluctuations. Reported times over 25 learning instances. Each data point in the box & whisker in **b)** is the median reported time for 100 recall trials (red bars in **a)**) for one particular learning instance. **c)** Evolution of weights in during training for network shown in a). Top, evolution of mean feed forward weights from column one to column two. Bottom, evolution of mean recurrent weights in column one.

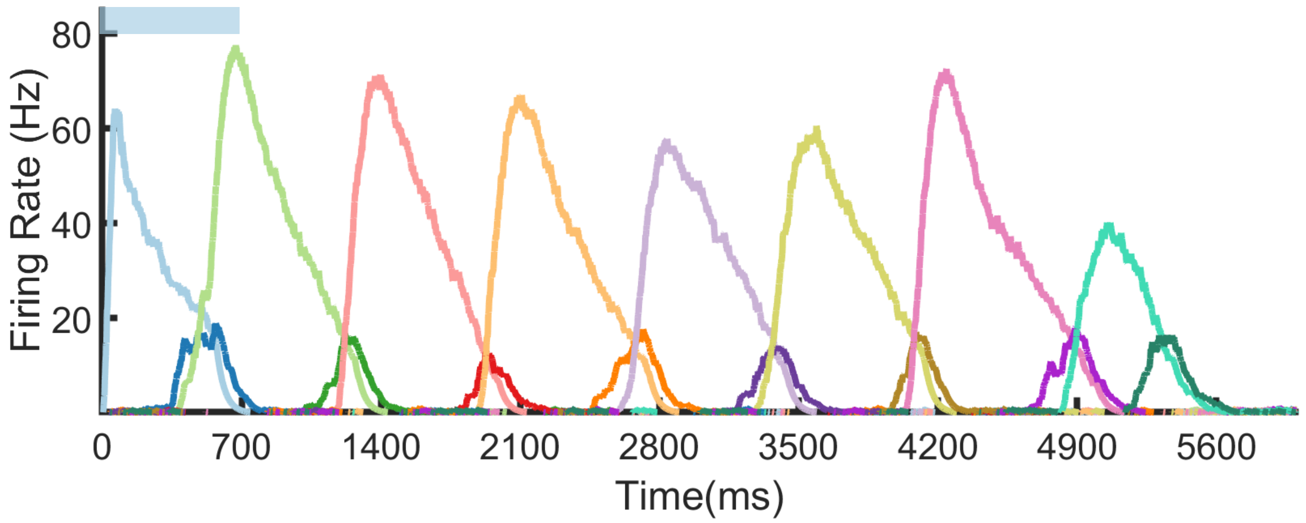


**Supplementary Figure 3. Eight Element Sequence Recall.** Recall after learning a sequence of eight elements, each with duration 700ms. Only the first element is stimulated. Notice that because of stochasticity, some elements (1 and 8) underreport their duration, while others (element 7) overreport their duration. In general, these errors can propagate in recall. For example, even though element 2 reports the correct duration (~700ms), the sequence still underreports 1400ms because of the errors in element 1.

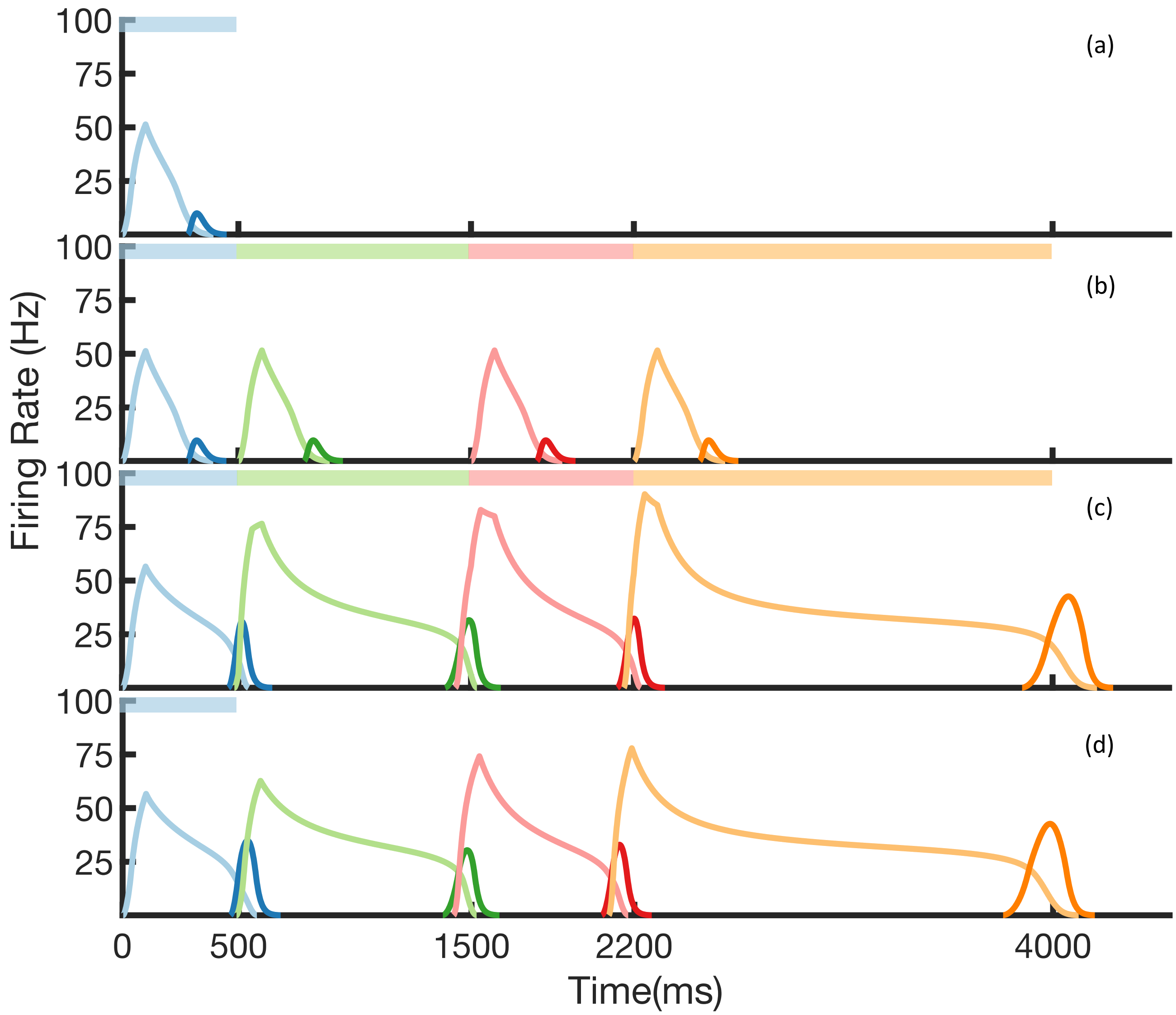


**Supplementary Figure 4. Rate-based learning and recall.** Recreation of Figure 3 from the main text, but using the rate-based formulation described in Methods. Each population of previously spiking neurons (e.g. red Timers) is now represented by one rate neuron. a) - d) Firing rates for timer cells (light colors) and messenger cells (dark colors) of 4 different columns during different stages of learning. Stimuli presented are shown as color bars in the top of plots. During learning, columns are stimulated in the sequence indicated by the colorbars (500, 1000, 700, and 1800 ms for blue, red, green, and orange, respectively). a) Before learning, the stimulation of a particular column only causes that column to be transiently active. b) During the first trial of learning, all columns in the sequence become activated by the stimuli, but have not yet learned to represent duration (through recurrent learning of Timer cells) or order (through feed forward learning of the Messenger cells). c) After many trials, the network learns to match the duration and order of presented stimuli. d) After learning, presenting the first element in the sequence is sufficient for recall of the entire sequence.

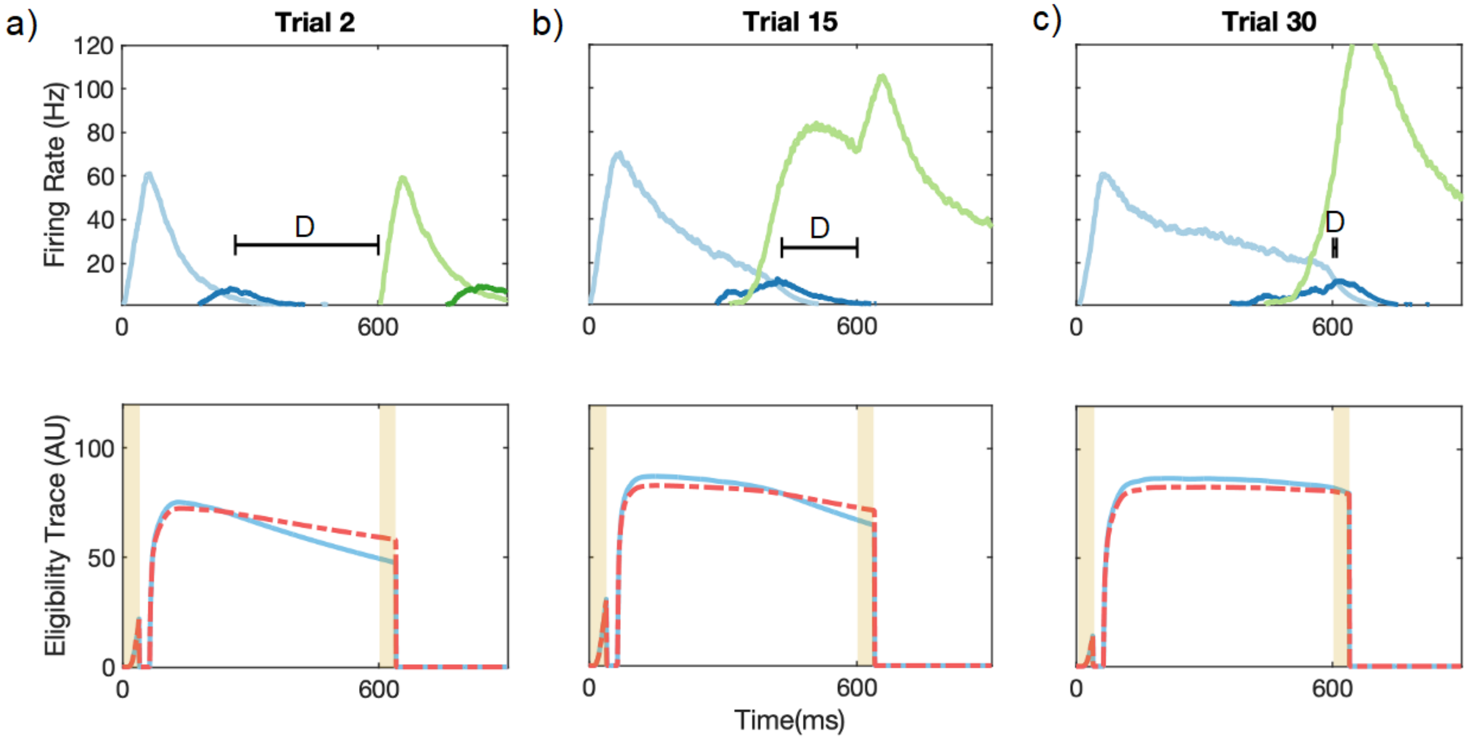


**Supplementary Figure 5. Recurrent Learning Evolution.** Top row: firing rates of timer (light colors) and messenger (dark colors) populations for the first two columns over the course of learning. Bottom row: eligibility traces corresponding to the recurrent weights for the timer cells in the first column (light blue in top row). **a)** For initial trials, LTP (dashed red line) dominates LTD (light blue line) in the reward windows (vertical yellow lines). This leads in a net increase in synaptic efficacy. Learning aims to minimize D, the time between the end of firing in one column and the beginning of firing in the next. **b)** For intermediate trials, the net difference (LTP-LTD) in the reward windows is still positive, but smaller than before. Synaptic efficacy continues to increase, but at a slower rate. **c)** As the timer population “extends” to represent the appropriate time interval (top), the net difference in the traces during the reward windows goes to zero. The synaptic weights reach a fixed point and learning is complete.

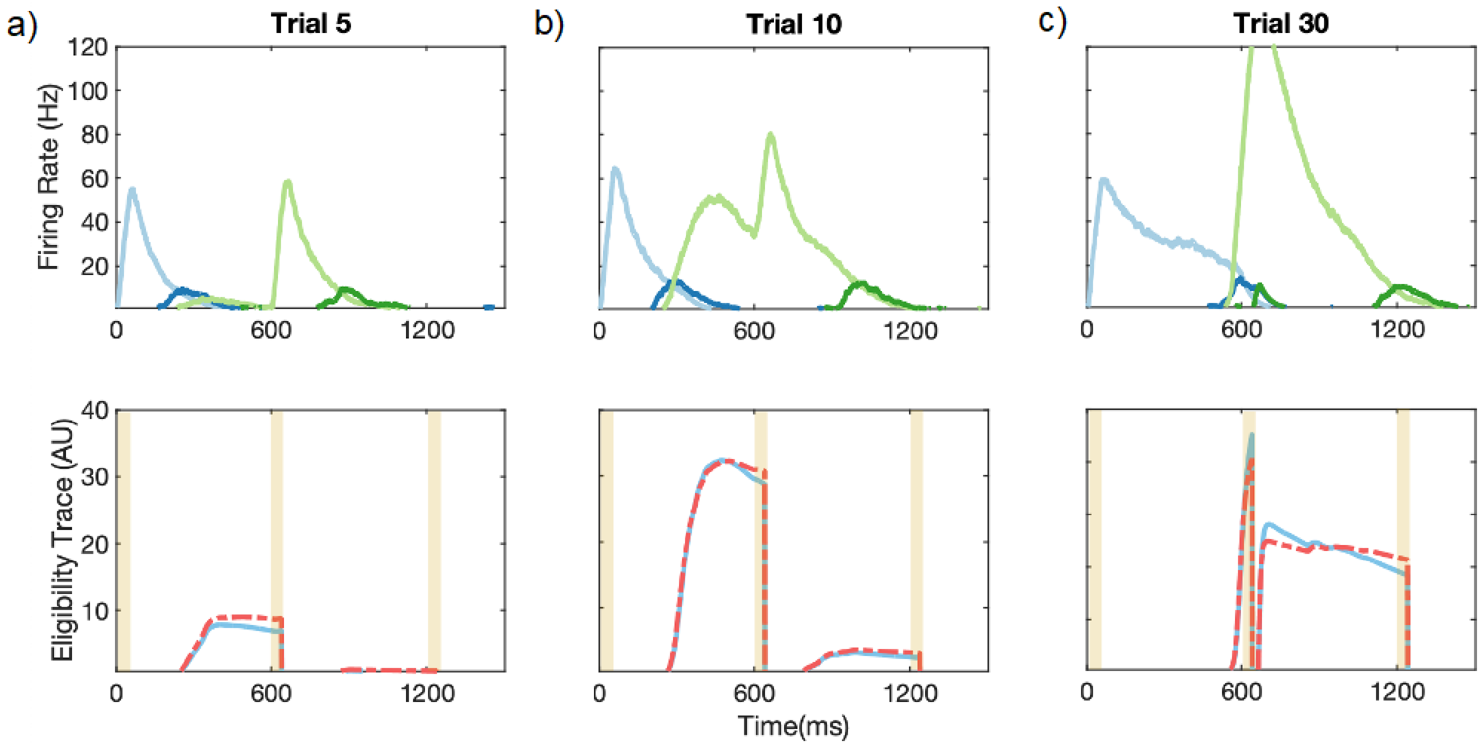


**Supplementary Figure 6. Feed Forward Learning Evolution.** Top row: firing rates of timer (light colors) and messenger (dark colors) populations for the first two columns. Bottom row: eligibility traces corresponding to the feed forward weights between the messenger cells of the first column (dark blue in top row) and the timer cells of the second column (light green in top row). **a)** For initial trials, LTP (dashed red line) dominates LTD (light blue line) in the reward windows (vertical yellow lines). This leads in a net increase in synaptic efficacy. **b)** For intermediate trials, the net difference (LTP-LTD) in the reward windows is still positive, but smaller than before. Synaptic efficacy continues to increase, but at a slower rate. **c)** As the messenger cells of the first column excite the timer cells of the second column to have an elevated firing rate (top), the net difference in the traces during the reward windows goes to zero. The synaptic weights reach a fixed point and learning is complete.

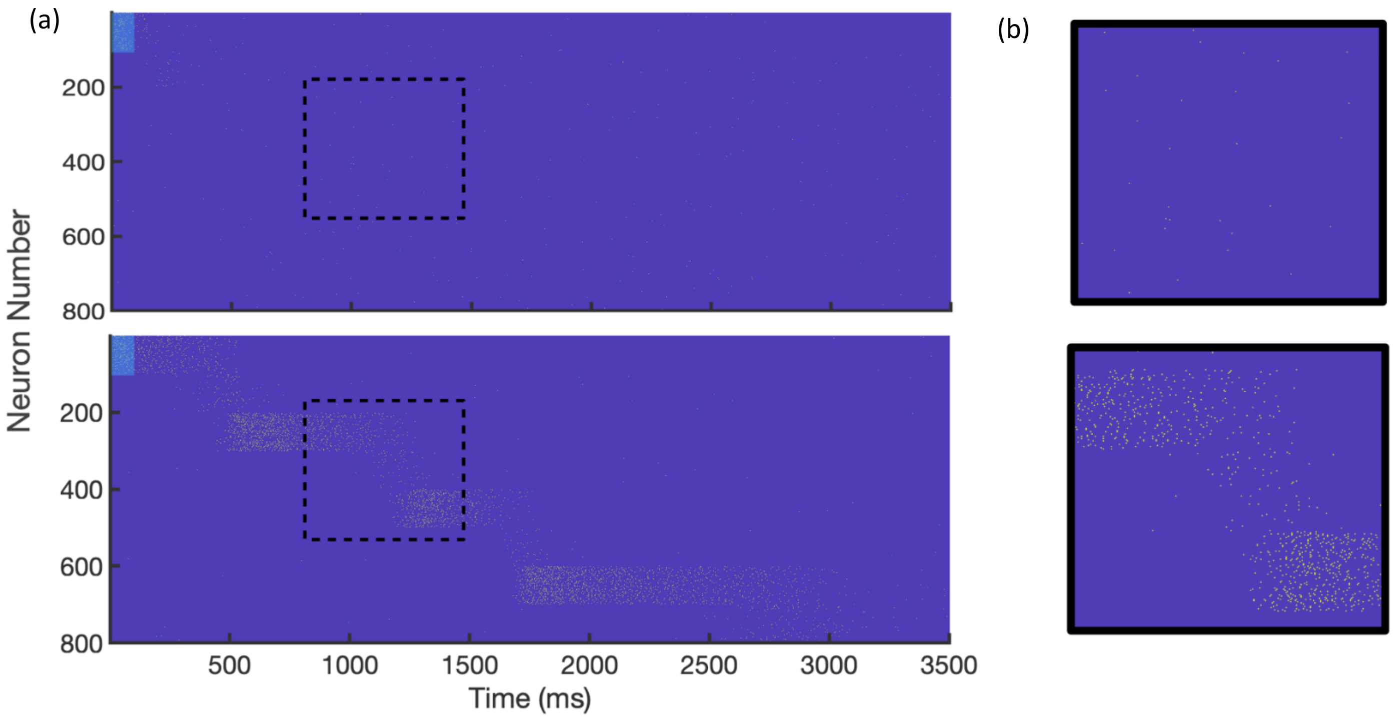


**Supplementary Figure 7. Spiking in learning network.** a) Spike rasters of network response to stimulation of first column (light blue bar) before (top), and after (bottom) learning a sequence of stimuli (500, 750, 500, and 1250 ms for columns 1, 2, 3, and 4, respectively). Neurons are sorted by population and sequentially by column. Neurons 1-100 are the Timer cells of the first column, neurons 101-200 are the Messenger cells of the first column, neurons 201-300 are the Timer cells of the second column, etc. b) Zoomed in insets of dashed boxes in a). Top, before learning, bottom, after learning.

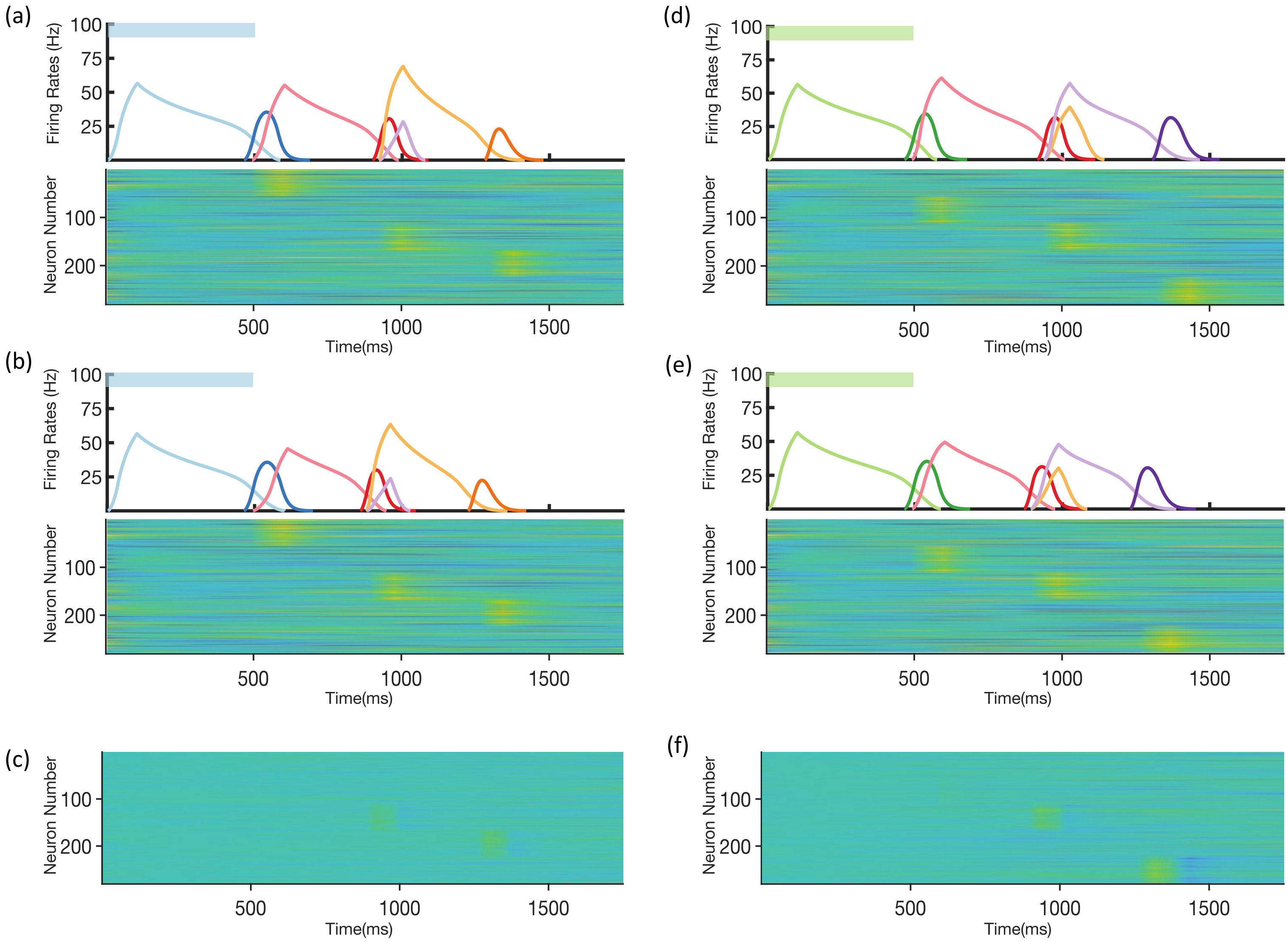


**Supplementary Figure 8. Robustness in non-Markovian recall.** A three-stage network trained on two non-Markovian sequences (BRO and GRP) recalls the two sequences with and without a perturbation to the initial state of the reservoir. a) The trained network is excited with a 500ms blue stimulus without an initial perturbation to the reservoir. Top, mean firing rates of Timer cells (light colors) and Messenger cells (dark colors) in the columnar network. Bottom, firing rates in the reservoir. The initial state of each unit in the reservoir is drawn from a normal distribution N(0,1.2), and the dynamics follow the equations in Methods. b) Same as in a) but a perturbation (normal distribution N(0,.25)) is applied to the initial state of the reservoir. In this example a slight sequence compression is observed, but the integrity of the transitions is maintained. c) The difference in reservoir firing rates between the perturbed trial and the unperturbed trial (same scale as a) and b)). d-f) Same as a-c), but this time stimulating with a 500ms green stimulus, as to recall sequence GRP.

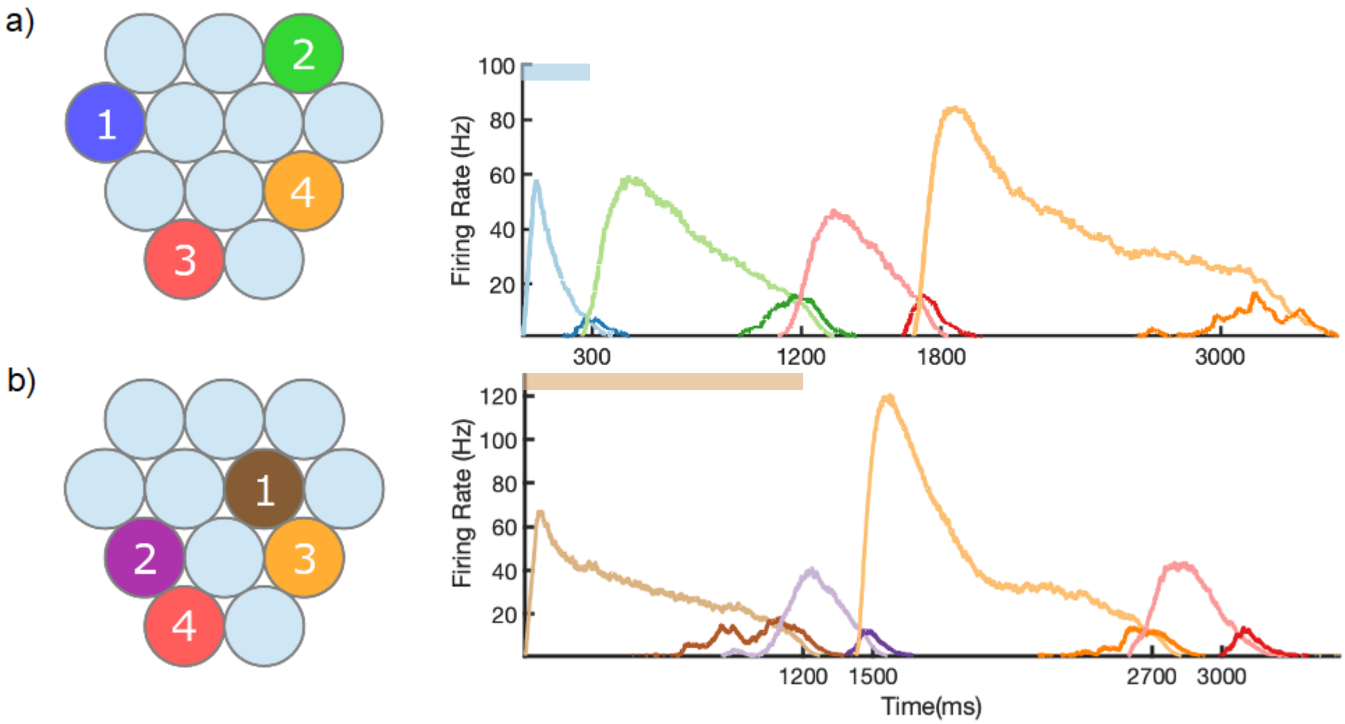


**Supplementary Figure 9. Learning of Different Sequences.** Left, identity and order of stimuli shown during training. Right, mean firing rate of network after training, upon stimulation of first column in sequence. a) Blue, green, red, and orange columns (numbers 4, 3, 11, 10) stimulated at times of 0, 300, 1200, and 1800 ms, respectively. b) Brown, purple, orange, and red columns (numbers 6, 8, 10, 11) stimulated at times of 0, 1200, 1500, and 2700, respectively.

| Parameter | Value | Units | Description |
|---|---|---|---|
| $N$ | 100 | - | Number of neurons per population |
| $dt$ | 1 | ms | Integration time step |
| $T$ | 50 | ms | Stimulus pulse duration |
| $\tau_{stim}$ | 50 | ms | Decay constant of stimulus |
| $\tau_w$ | 40 | ms | Time window for firing rate integration |
| $p_r$ | 40 | Hz | Rate of Poisson stimulus pulse |
| $\rho$ | 1/7 | - | Fractional change of synaptic activation |
| $\tau_s^E$, $\tau_s^I$ | 80,10 | ms | Time constant for synaptic activation for excitatory (EE) and inhibitory (EI, IE) connections |
| $g_L$ | .001 | μS | Leak conductance |
| $C_m$ | 20 x $g_L$ | nF | Membrane capacitance |
| $E_L$ | -60 | mV | Leak reversal potential |
| $E_E$, $E_I$ | -5,-70 | mV | Excitatory and inhibitory reversal potentials |
| $v_{th}$ | -55 | mV | Spiking threshold potential |
| $v_{rest}$ | -60 | mV | Resting potential |
| $v_{hold}$ | -61 | mV | Reset potential |
| $t_{ref}$ | 2 | ms | Absolute refractory period |
| $\tau_p$, $\tau_d$ | 2000, 1000 | ms | LTP/LTD eligibility trace time constant |
| $T_p^{max}$, $T_d^{max}$ | 0.95, 1 | - | Saturation level, LTP/LTD eligibility trace |
| $\eta_p$, $\eta_d$ | 1, 0.55 | $ms^{-1}$ | Activation rate, LTP/LTD eligibility trace |
| $\tau_p^{FF}$, $\tau_d^{FF}$ | 200, 800 | ms | LTP/LTD eligibility trace time constant, feed forward connections |
| $T_p^{max,FF}$, $T_d^{max,FF}$ | 0.98, 1 | - | Saturation level, LTP/LTD eligibility trace, feed forward connections |
| $\eta_p^{FF}$, $\eta_d^{FF}$ | 0.44, 0.33 | $ms^{-1}$ | Activation rate, LTP/LTD eligibility trace, feed forward connections |
| $T_{reward}$ | 25 | ms | Duration of neuromodulator presentation upon change in stimulus |
| $T_{tr}$ | 25 | ms | Duration of refractory period for traces following neuromodulator presentation |
| $\eta_{rec}$, $\eta_{FF}$ | .0045, .08 | $ms^{-1}$ | Learning rates, recurrent and feed forward connections |
| $\varphi$ | 0.3 | - | Sparsity of fixed connections |
| $W_{EE}^{MT}$, $W_{EI}^{MT}$ | .02, .7 | μS | Synaptic connection strength, Timer to Messenger excitatory to excitatory (EE) and inhibitory to excitatory (EI) connections |
| $W_{EI}^{TT}$, $W_{EI}^{MM}$ | 1, 1 | μS | Synaptic connection strength, intercolumnar Timer-Timer and Messenger-Messenger inhibitory to excitatory (EI) connections |
| $W_{IE}^{TT}$, $W_{IE}^{MM}$ | .002, .01 | μS | Synaptic connection strength, intracolumnar Timer-Timer and Messenger-Messenger excitatory to inhibitory (IE) connections |

**Supplementary Table 1. List of Main Model Parameters.**

| Parameter | Value | Units | Description |
| --- | --- | --- | --- |
| $K$ | 280 | - | Number of units in reservoir |
| $g$ | 1.5 | μS | Gain parameter |
| $M$ | 1120 | - | Number of units in sparse pattern net |
| $\boldsymbol{\theta_m}$ | 7.5 | Hz | Threshold for columnar to reservoir excitation |
| $\boldsymbol{\theta_o}$ | 0.1 | - | Threshold for reservoir to sparse net excitation |
| $\mathbf{Q_{max}}$ | .15 | μS | Rate of Poisson stimulus pulse |
| $\boldsymbol{\tau_{net}}$ | 100 | ms | Time constant for units in reservoir |
| $\boldsymbol{\tau_u}, \boldsymbol{\tau_{ui}}$ | 40,10 | ms | Excitatory and inhibitory time constants, rate-based network |
| $\mathbf{u_c}$ | 2 | - | Upper threshold, rate-based transfer function |
| $\boldsymbol{\theta}$ | 0 | - | Lower threshold, rate-based transfer function |
| $\boldsymbol{\nu}$ | 2 | - | Scaling parameter, rate-based transfer function |

**Supplementary Table 2. List of Reservoir, Sparse Net, and Rate-Based Model Parameters.**
